# UPF1 shuttles between nucleus and cytoplasm independently of its RNA binding and ATPase activities

**DOI:** 10.1101/2025.03.24.645002

**Authors:** Sofia Nasif, Andrea Brigitte Eberle, Karin Schranz, Remo Hadorn, Sutapa Chakrabarti, Oliver Mühlemann

## Abstract

The ATP-dependent RNA helicase Up-frameshift 1 (UPF1) is an essential protein in mammalian cells and a key factor in nonsense-mediated mRNA decay (NMD), a translation-dependent mRNA surveillance process. UPF1 is mainly cytoplasmic at steady state but accumulates in the nucleus after inhibiting CRM1-mediated nuclear export by Leptomycin B (LMB), indicating that UPF1 shuttles between the nucleus and the cytoplasm. Consistent with its dual localization, there is evidence for nuclear functions of UPF1, for instance in DNA replication, DNA damage response, and telomere maintenance. However, whether any of UPF1’s biochemical activities are required for its nuclear-cytoplasmic shuttling remains unclear.

To investigate this, we examined two UPF1 mutants: the well-described ATPase-deficient UPF1-DE (D636A/E637A) and a newly generated RNA-binding mutant UPF1-NKR (N524A/K547A/R843A). Biochemical assays confirmed that the UPF1-NKR mutant cannot bind RNA or hydrolyze ATP *in vitro* but retains interaction with UPF2, UPF3B, and SMG6. Overexpression of UPF1-NKR exerted a dominant-negative effect on endogenous UPF1 and inhibited NMD. Subcellular localization studies revealed that UPF1-DE accumulates in cytoplasmic granules (P-bodies), even in the presence of LMB, whereas UPF1-NKR shuttles normally. This indicates that UPF1’s shuttling does not require its RNA-binding or ATPase activities. Notably, the UPF1-DE.NKR double mutant restored nuclear-cytoplasmic shuttling and prevented accumulation in P-bodies, suggesting that the shuttling defect of UPF1-DE arises from its tight binding to RNA. Overall, our findings demonstrate that UPF1’s shuttling is independent of its ATPase and RNA-binding activities, with RNA binding itself being a key determinant of its cytoplasmic retention.

## Introduction

Up-frameshift 1 (UPF1) was described in the early 90s as a factor involved in the rapid decay of mRNAs containing premature translation termination codons (PTCs) in *S. cerevisiae* (Leeds et al. 1991) and *C. elegans* (Pulak and Anderson 1993). Shortly after its discovery, it was characterized as a nucleic acid helicase with nucleic acid-dependent ATPase activity belonging to the helicase superfamily 1 (SF1) (Czaplinski et al. 1995; Weng et al. 1996), and within the following decade, UPF1 homologs were described in other species like *H. sapiens* (Perlick et al. 1996; Applequist et al. 1997), *D. melanogaster* (Gatfield et al. 2003) and *A. thaliana* (Yoine et al. 2006). Since its discovery, UPF1 has been implicated in various cellular processes, with its role as the central regulator of nonsense-mediated mRNA decay (NMD) being the most extensively studied (Kim and Maquat 2019; Karousis and Muhlemann 2022).

NMD is a eukaryotic translation-dependent mRNA degradation pathway that serves both as a quality control mechanism, by targeting for degradation faulty mRNAs containing PTCs, as well as a post-transcriptional regulator of gene expression, by modulating the levels of a subset of mRNAs with full coding potential (Nasif et al. 2017; Kishor et al. 2019). The precise molecular mechanism of NMD is still incompletely understood due to its inherent heterogeneity. NMD exhibits context-dependent regulation, which varies across cellular environments and species and involves distinct trans-acting factors, resulting in mechanistic differences. (Nasif et al. 2017; Lloyd 2018; Sato and Singer 2021; Karousis and Muhlemann 2022). For example, different species harbor distinct repertoires of NMD factors, with only UPF1 being present in all eukaryotes, including the early branching Excavates (Lloyd 2018). Although UPF1 plays a central and undisputed role in NMD, the mechanisms by which it identifies and selects its targets are still not fully understood. UPF1 binds without apparent sequence specificity to mRNAs through its conserved helicase core composed of two RecA-like domains that form a composite RNA-binding surface and an ATP-binding pocket. The helicase core of UPF1 is N-terminally flanked by a cysteine and histidine-rich domain (CH), and additionally contains flexible domains rich in phosphorylatable serine-glutamine and threonine-glutamine ([S/T]Q) motifs in its N-and C-termini (Cheng et al. 2007; Chakrabarti et al. 2011). Driven by ATP hydrolysis, UPF1 can translocate 5‘ to 3’ along nucleic acids, displace proteins bound to them, and dissociate from RNAs (Czaplinski et al. 1995; Weng et al. 1996; Fiorini et al. 2015; Chapman et al. 2022). Both the N-and C-terminal domains were shown to suppress UPF1’s catalytic activities *in vitro* (Chamieh et al. 2008; Fiorini et al. 2013). UPF1’s interaction partner UPF2 can bind to the CH domain, and by inducing an extensive conformational change in UPF1, UPF2 relieves the autoinhibitory function and enhances UPF1’s catalytic activities (Chamieh et al. 2008; Chakrabarti et al. 2011; Chapman et al. 2024). In addition, phosphorylated N-and C-terminal [S/T]Q motifs serve as binding platforms to recruit downstream NMD factors that trigger the decay of the mRNA (Huntzinger et al. 2008; Eberle et al. 2009; Okada-Katsuhata et al. 2012; Durand et al. 2016). The ability of UPF1 to dissociate from non-NMD-targeted mRNAs through ATP hydrolysis has been proposed as a key determinant of its substrate specificity, as ATPase-deficient UPF1 mutants exhibit indiscriminate binding to both NMD and non-NMD substrates (Kurosaki et al. 2014; Lee et al. 2015). Furthermore, ATPase-driven RNA dissociation by UPF1 is essential for the completion of mRNA degradation, as ATPase-deficient UPF1 mutants remain bound to decay intermediates and block the access of exonucleolytic enzymes (Franks et al. 2010).

With approximately 450,000 molecules per cell, mammalian UPF1 is an abundant protein that is predominantly localized in the cytoplasm under physiological conditions, consistent with its role in promoting NMD as well as other translation-dependent RNA decay pathways (Atkin et al. 1995; Applequist et al. 1997; Lykke-Andersen et al. 2000; Mendell et al. 2002; Franks et al. 2010; Imamachi et al. 2012; Kim and Maquat 2019; Cho et al. 2022). UPF1 has also been shown to localize to the endoplasmic reticulum (ER), in an NBAS-dependent manner, to aid in the degradation of NMD substrates that are translated at the ER (Longman et al. 2020). Additionally, experimental evidence suggests that a fraction of UPF1 shuttles between the cytoplasm and the nucleus. For instance, treatment with leptomycin B (LMB), a *Streptomyces* metabolite that blocks CRM1-dependent nuclear export (Kudo et al. 1998), results in nuclear accumulation of UPF1 in both human and *Drosophila* cells (Mendell et al. 2002; Singh et al. 2019). Moreover, UPF1 has been detected associated with chromatin fractions after cellular fractionation and in chromatin immunoprecipitation experiments (Azzalin and Lingner 2006b; Azzalin et al. 2007; Chawla et al. 2011; Singh et al. 2019). Furthermore, UPF1 has been shown to interact with nuclear proteins like the telomeric factor TPP1, telomerase (Chawla et al. 2011), as well as with subunits of the DNA polymerase δ (Carastro et al. 2002; Azzalin and Lingner 2006b). In accordance with its nuclear-cytoplasmic shuttling, also nuclear functions have been reported for UPF1. For instance, it has been suggested that UPF1 is involved in nonsense-mediated altered splicing (Mendell et al. 2002), DNA replication and cell cycle progression (Azzalin and Lingner 2006b), telomere homeostasis (Azzalin et al. 2007; Chawla et al. 2011), and the DNA damage response (Azzalin and Lingner 2006b). In *Drosophila*, it has also been reported that UPF1 is associated with active Pol II transcription sites and that it is required for the efficient release of mRNAs from chromatin and their export to the cytoplasm (Singh et al. 2019). Similarly, UPF1 has been implicated in the nuclear export of HIV-1 genomic RNA (Ajamian et al. 2015). UPF1’s capacity to bind DNA and RNA, its ability to use ATP hydrolysis to translocate along nucleic acids and remodel them, and its shuttling nature make it a strong candidate for the proposed nuclear functions. However, to unambiguously distinguish UPF1’s nuclear and cytoplasmic functions, a better understanding of the nuclear import and export pathways used by UPF1, as well as how its shuttling is affected by cellular conditions and its own biochemical activities, is required.

In this study, we investigated if UPF1’s biochemical activities are determinants of its subcellular localization. To this end, we designed and characterized a novel RNA-binding mutant, UPF1-NKR, by introducing three point mutations (N524A, K547A, R843A). We confirmed that this mutant does not bind RNA *in vitro* and consequently fails to hydrolyze ATP. Furthermore, UPF1-NKR is unable to restore NMD activity in UPF1-depleted cells and exerts a dominant-negative effect on endogenous UPF1 when overexpressed in mammalian cells. By analyzing the subcellular distribution of various UPF1 mutants, we demonstrate that UPF1’s nuclear-cytoplasmic shuttling is independent of its ATPase and RNA binding activities.

## Results

### The UPF1-NKR mutant protein fails to bind RNA and hydrolyze ATP *in vitro*

To test if RNA binding is important for UPF1’s ability to shuttle between the nucleus and the cytoplasm, we designed point mutations to abolish UPF1 binding to RNA. Based on the structure of yeast Upf1 (yUpf1) bound to RNA, we identified amino acids asparagine 462, lysine 485, and arginine 779 as key residues that interact with RNA. Based on the high sequence and structural homology of yeast and human UPF1, we predicted that the homologous residues in human UPF1, amino acids asparagine 524, lysine 547, and arginine 843 would be engaged in and therefore presumably be required for RNA binding. We mutated these residues to alanines to generate the RNA-binding mutant UPF1-NKR (Fig. 1). The ability of this mutant to bind RNA *in vitro* was tested using MicroScale Thermophoresis (MST). To this end, we expressed and affinity-purified C-terminally 3xFlag-tagged full-length human UPF1 in three different versions: wild type (WT), the RNA binding mutant (NKR), and the well-characterized ATPase-deficient D636A/E637A (DE) (Weng et al. 1996; Franks et al. 2010), as well as maltose-binding protein (MBP) as a control protein, from Flp-In T-Rex 293 cells (Fig. 2A). To measure RNA binding by MST, the purified recombinant proteins were titrated (300 – 0.009 nM) against a constant amount (4 nM) of Cy5-labeled U30 RNA, which was the lowest concentration of the labeled RNA molecule we could confidently detect with this assay. We first did a time-course experiment to determine the time required to reach the binding equilibrium between UPF1-WT and the U30 RNA and found this to be 60 min at 25 °C (Suppl. Fig. 1A). Similar experiments performed with the UPF1-DE protein showed that this mutant reaches binding saturation faster than the UPF1-WT counterpart, probably due to its irreversible binding to RNA (Suppl. Fig. 1A). We also addressed the impact of ATP in the binding reactions, as it was shown that ATP binding and hydrolysis reduce the affinity of UPF1 for nucleic acids (Dehghani-Tafti and Sanders 2017; Gowravaram et al. 2018; Chapman et al. 2022). We therefore performed the binding reactions for 60 min at 25 °C, in the presence or absence of 1 mM ATP, followed by MST to assess the RNA binding activities of all the purified recombinant proteins (Fig. 2B). As predicted, the UPF1-NKR mutant did not bind RNA *in vitro*, showing comparable MST traces as MBP, regardless of ATP presence (Fig. 2B). In contrast, UPF1-WT and UPF1-DE proteins bound RNA with characteristic sigmoidal dose-response curves, allowing estimation of dissociation constants (*K_d_*). We found that in the absence of ATP, UPF1-WT and UPF1-DE proteins bound to RNA with similar affinities, with calculated *K_d_* of 16.378 ± 0.871 nM and 25.731 ± 1.277 nM, respectively (Fig. 2B). While the presence of ATP in the reaction did not affect the RNA binding affinity of the UPF1-DE protein (calculated *K_d_* 17.697 ± 1.095 nM), we observed a 50-fold reduction in the RNA binding capacity of the UPF1-WT protein with an estimated *K_d_* of 873.5 ± 305.1 nM. These results are in agreement with previous reports showing that ATP binding and hydrolysis reduce the affinity of UPF1-WT for RNA (Dehghani-Tafti and Sanders 2017; Gowravaram et al. 2018; Chapman et al. 2022). However, under our experimental conditions, the presence of ATP resulted in a larger *K_d_* reduction than the previously reported 4 to 18-fold changes (Dehghani-Tafti and Sanders 2017; Gowravaram et al. 2018; Chapman et al. 2022).

**Figure 1.**
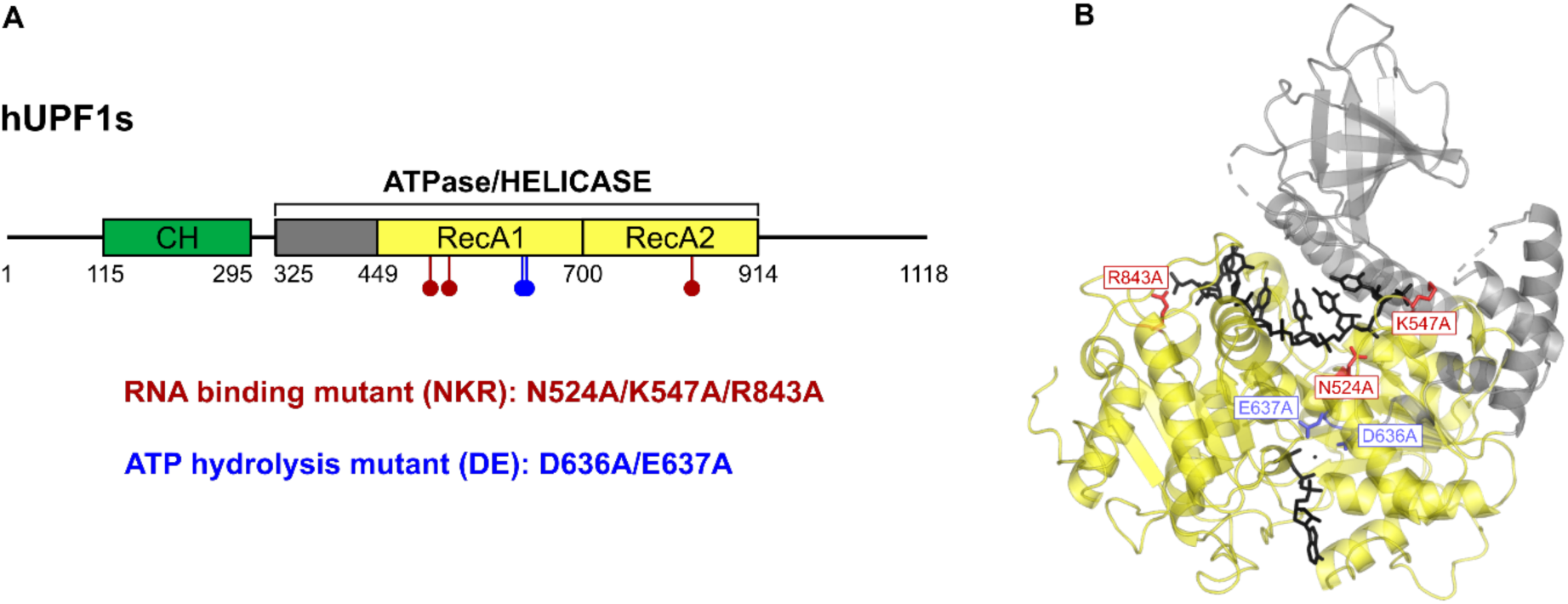
Design of the RNA binding mutant UPF1-NKR. (A) Scheme of the human UPF1 short isoform, with the amino acid changes introduced in the RNA binding mutant (NKR) depicted in red or the ATPase mutant (DE), shown in blue. (B) The localization of the NKR and DE residues is depicted within the crystal structure of human Upf1ΔCH in complex with six ribonucleotides of a U_15_ RNA and ADP:AlF_4_^−^.

**Figure 2.**
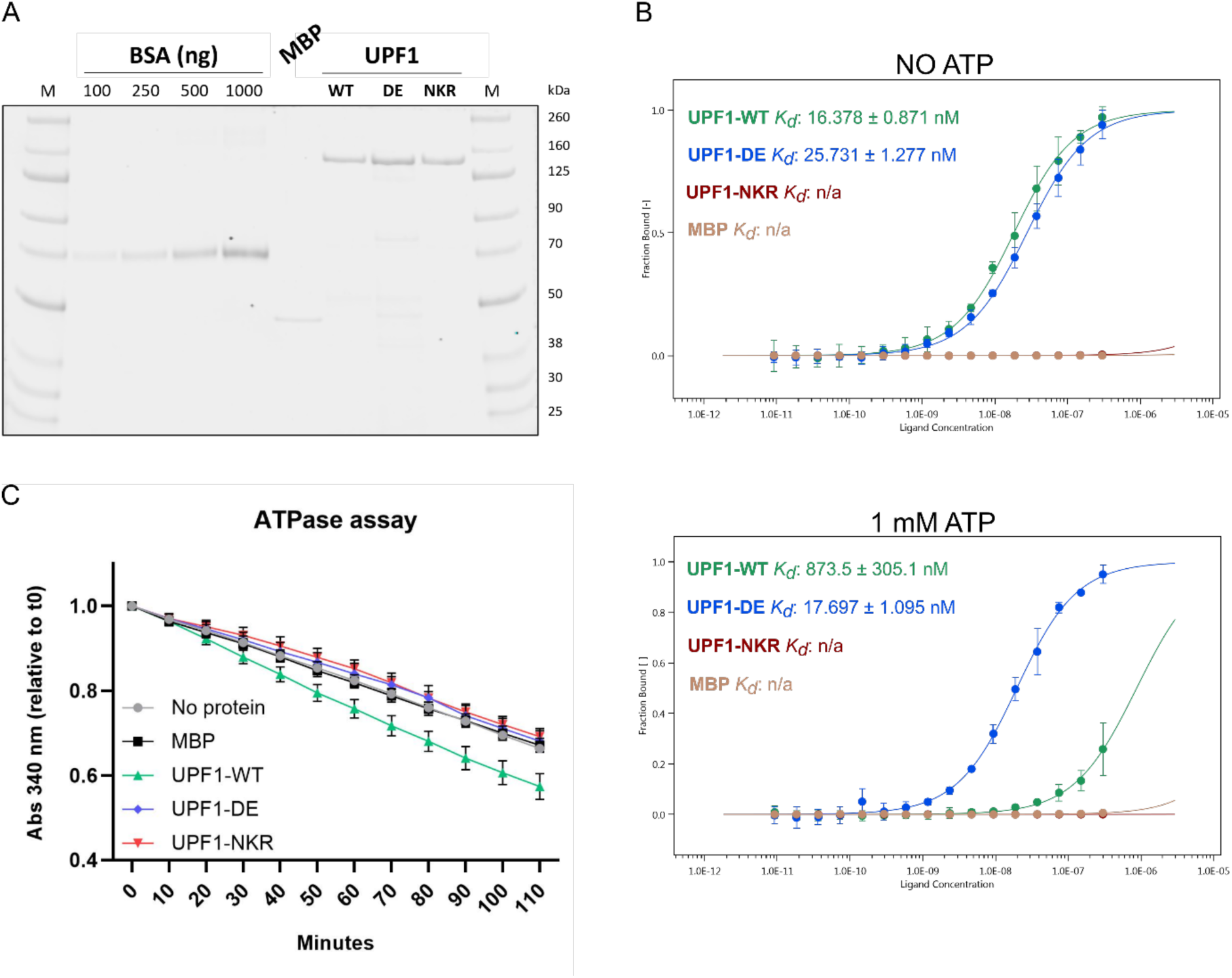
The UPF1-NKR mutant protein fails to bind RNA and hydrolyze ATP *in vitro*. (A) Coomassie-stained gel showing the purified UPF1-3xFlag WT, DE, and NKR mutants and MBP-3xFlag purified as a control. A BSA standard curve loaded on the same gel was used to estimate the concentration of the purified proteins. M = molecular weight marker. (B) *In vitro* RNA binding was analyzed by MicroScale Thermophoresis (MST). Binding reactions were incubated for 1 h at 25 °C in the absence or presence of ATP, followed by MST. The fraction of bound RNA is plotted against the final concentration of the unlabeled titrated proteins, and the averages and standard deviations from three independent dilution series of UPF1 are shown. Estimated dissociation constants (*K_d_*) are shown for UPF1-WT and UPF1-DE proteins; n/a = not available. (C) NADH-coupled ATPase assays were performed for 2 h at 37 °C, the oxidation of NADH was followed by measuring absorbance at 340 nm. Shown are the averages and standard deviations of 4-6 independent measurements.

Since UPF1 shows RNA-dependent ATPase activity (Czaplinski et al. 1995; Bhattacharya et al. 2000; Cheng et al. 2007; Chapman et al. 2022), we predicted that the UPF1-NKR mutant would be unable to hydrolyze ATP *in vitro*, and we used NADH-coupled ATPase assays to test our hypothesis. We first compared the *in vitro* ATPase activity of the full-length UPF1-WT purified from mammalian cells to that of the UPF1 helicase domain (UPF1-HD) expressed in *E. coli*. We observed that both proteins perform RNA-dependent ATP hydrolysis, with the UPF1-HD showing a slightly higher activity than the full-length UPF1-WT protein, presumably due to the presence of the auto-inhibitory N-and C-terminal domains in full-length UPF1 (Chamieh et al. 2008; Chakrabarti et al. 2011; Fiorini et al. 2013) (Suppl. Fig. 1B). Consistent with our prediction, the UPF1-NKR mutant failed to hydrolyze ATP *in vitro*, as did the UPF1-DE mutant and the negative control MBP (Fig. 2C). Altogether, our results show that the UPF1-NKR mutant is indeed unable to bind RNA and, consequently, also fails to hydrolyze ATP *in vitro*.

### UPF1-NKR mutant is not functional in NMD

Since the UPF1-NKR protein is unable to bind RNA and hydrolyze ATP, we expected that this mutant would also fail to sustain NMD activity in mammalian cells. To test this, we performed knockdown (KD) and rescue experiments by stably integrating pKK plasmids into Flp-In T-Rex 293 cells (Szczesny et al. 2018). These plasmids harbor a Doxycycline (Dox)-inducible bidirectional promoter that allows the simultaneous expression of an RNA interference cassette (RNAi) for depletion of an endogenous protein in one direction and an RNAi-resistant recombinant version of the targeted protein in the other direction. We established cell lines where endogenous UPF1 can be depleted and replaced with either MBP, UPF1-WT, UPF1-DE, or UPF1-NKR proteins, each carrying a 3xFlag tag in their C-terminus. As controls, we generated cell lines that express a luciferase-targeting RNAi (CTRL KD) and the MBP protein. To optimize the KD and rescue conditions, we analyzed NMD activity in cells at various time points following Dox induction using RT-qPCR. Under UPF1 KD conditions, we observed a time-dependent accumulation of all three tested NMD-sensitive transcripts. The RP9P transcript reached its maximum levels after 72 hours of Dox induction, whereas the hnRNPL_NMD and TRA2B_NMD transcripts showed the highest accumulation after 96 hours of induction (Suppl. Fig. 2). NMD activity could be efficiently rescued by the expression of UPF1-WT for all tested time points. The observed accumulation of transcripts is specific for NMD targets since an NMD-insensitive transcript like the hnRNPL protein-coding splice isoform (hnRNPL_Prot) remained unchanged throughout the different time points and experimental conditions (Suppl. Fig. 2). Next, we compared the NMD activity of all the generated cell lines after 96 h of Dox induction (Fig. 3). We observed the specific accumulation of the NMD-sensitive transcripts when the endogenous UPF1 protein was depleted and the unrelated control protein MBP was expressed (UPF1 KD + MBP, Fig. 3). As expected, the expression of the UPF1-WT protein resulted in a complete rescue of NMD activity and restored the levels of the NMD-sensitive transcripts to that of the control condition (UPF1 KD + UPF1-WT, Fig. 3). On the contrary, expression of the UPF1-DE or the UPF1-NKR mutants failed to restore NMD activity, results that are consistent with UPF1’s ATPase and RNA binding activities being essential for NMD (UPF1 KD + UPF1-DE and UPF1 KD + UPF1-NKR, Fig. 3). We also observed that overexpression of UPF1-WT protein in addition to the endogenous UPF1 does not result in increased NMD activity, as evidenced by the unaffected levels of the NMD-sensitive transcripts in the CTRL KD + UPF1-WT condition (Fig. 3). These results suggest that UPF1 protein levels are not rate-limiting for NMD in Flp-In T-Rex 293 cells, as was already shown for HeLa cells (Huang et al. 2011). The analysis of these cell lines confirmed that the UPF1-NKR mutant cannot sustain NMD activity.

**Figure 3.**
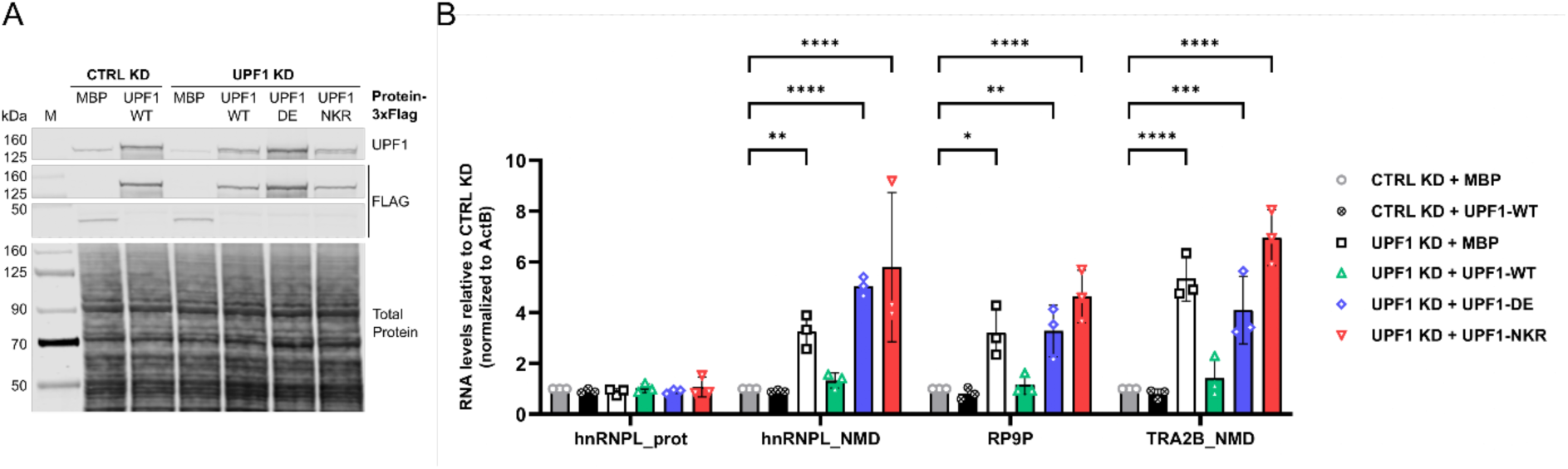
UPF1-NKR mutant is not functional in NMD. HEK cells stably transfected with pKK plasmids were induced for 96 h with Doxycycline to simultaneously express siRNAs against UPF1 (UPF1 KD) or luciferase (CTRL KD), and RNAi-resistant UPF1 versions (WT, DE, or NKR) or MBP as a control. (A) The efficiency of the UPF1 KD and the expression levels of the recombinant proteins were assessed by western blotting with the indicated antibodies. Total protein stain served as loading control. M = molecular weight marker. (B) Relative mRNA levels of hnRNPL_Protein, hnRNPL_NMD, RP9P, and TRA2B NMD, normalized to actin mRNA levels, were determined by RT-qPCR. Averages and standard deviations from three independent experiments are shown. Statistical significance was determined by Two-way ANOVA and Dunnett’s multiple comparisons test against the CTRL KD + MBP condition. Asterisks are only displayed for the comparisons with a *p*-value smaller than 0.05. * p ≤ 0.05; ** p ≤ 0.01; *** p ≤ 0.001; **** p ≤ 0.0001.

### UPF1’s subcellular localization is determined independently of its RNA binding or ATPase activities

Consistent with its central role in NMD, UPF1 is mainly cytoplasmic at steady state in mammalian cells. However, UPF1 can shuttle between the nucleus and the cytoplasm, as evidenced by its nuclear accumulation after Leptomycin B (LMB) treatment, which blocks CRM1-dependent nuclear export (Kudo et al. 1998; Mendell et al. 2002; Singh et al. 2019). Studies in *Drosophila* have shown that UPF1 is continuously shuttling between the nucleus and the cytoplasm, and the fact that the UPF1-DE mutant fails to accumulate in the nucleus after LMB treatment, has led to the suggestion that UPF1 shuttling depends on its ATPase activity (Singh et al. 2019). Whether any of UPF1’s biochemical activities are required for its shuttling in mammalian cells has not been addressed. We therefore set out to investigate if UPF1’s RNA binding or ATPase activities are important in determining its subcellular localization in HeLa cells. For this, we overexpressed UPF1-WT, UPF1-DE, and UPF1-NKR proteins with a C-terminal GFP-tag, and studied their subcellular localization after LMB or ethanol (control) treatment (Fig. 4). Consistent with previous findings, the UPF1-WT-GFP protein can be detected exclusively in the cytoplasm in the ethanol-treated condition, and a fraction of it accumulates in the nucleoplasm after LMB treatment (Mendell et al. 2002; Singh et al. 2019). The localization of the overexpressed UPF1-DE-GFP protein also aligns with previous reports, showing a cytoplasmic localization under ethanol treatment and being barely detectable in the nucleus in LMB-treated cells, indicating that UPF1-DE lost its ability to shuttle (Mendell et al. 2002; Singh et al. 2019). Moreover, we also observe its accumulation in bright cytoplasmic dots, previously identified as processing bodies (P bodies) (Sheth and Parker 2006; Cheng et al. 2007; Stalder and Muhlemann 2009; Franks et al. 2010; Singh et al. 2019). Interestingly, the UPF1-NKR-GFP protein shows the same shuttling behavior as the UPF1-WT-GFP protein: cytoplasmic at steady state with a marked accumulation in the nucleoplasm upon LMB treatment (Fig. 4). Since the NKR mutant fails to bind RNA as well as to hydrolyze ATP, this result suggests that neither of these activities is responsible for determining UPF1’s subcellular localization. Given this finding, we hypothesized that the failure of the UPF1-DE-GFP protein to shuttle to the nucleus might stem from its tight binding to cytoplasmic RNAs, from which it cannot dissociate in the absence of ATPase activity. If this reasoning was correct, the addition of the NKR mutations in the background of the UPF1-DE mutant should restore the nuclear-cytoplasmic shuttling capacity of the resulting UPF1-DE.NKR double mutant. To test this prediction, we generated the UPF1-DE.NKR double mutant and confirmed that it fails to bind RNA *in vitro* (Suppl. Fig. 3). We next studied the subcellular localization of the UPF1-DE.NKR fused to GFP. Consistent with our hypothesis that the tight binding to RNA prevents shuttling of UPF1-DE, the UPF1-DE.NKR-GFP protein is cytoplasmic under ethanol treatment and accumulates in the nucleus upon LMB treatment (Fig. 4). Also, the localization of the UPF1-DE-GFP protein to P bodies is reverted in the UPF1-DE.NKR-GFP mutant, indicating that RNA binding is required for the localization of UPF1-DE to P bodies. Overall, our results demonstrate that UPF1’s nuclear-cytoplasmic shuttling is independent of its RNA binding or ATPase activities.

**Figure 4.**
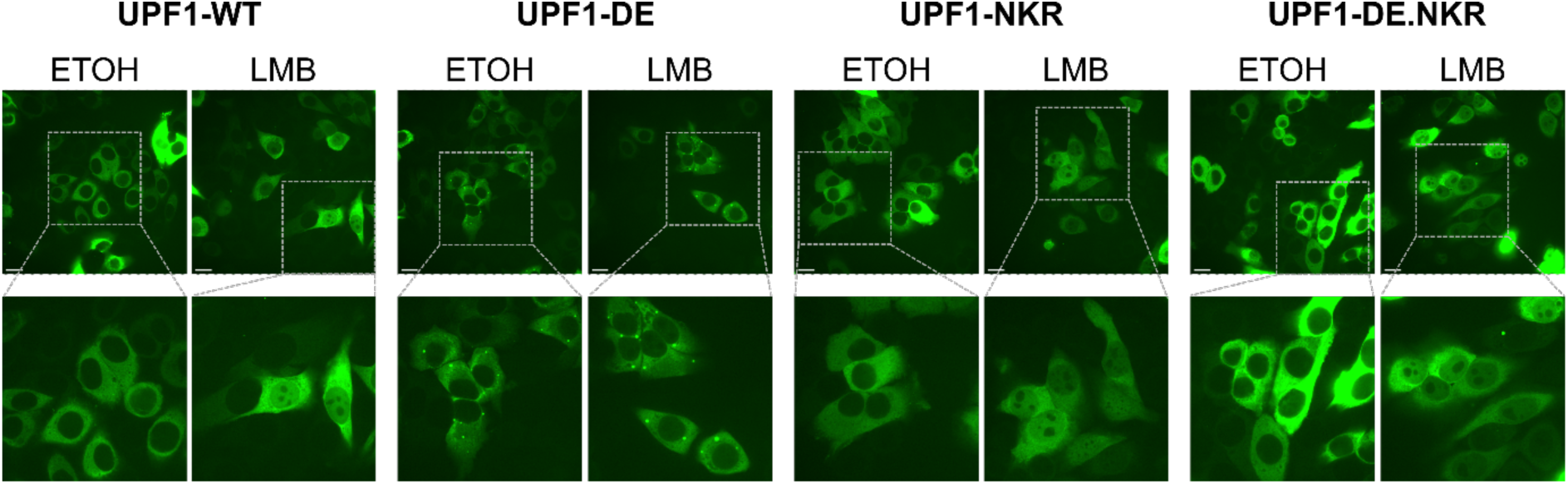
UPF1 shuttles between the nucleus and the cytoplasm independently of its RNA-binding and ATPase activities. UPF1-GFP (WT, DE, NKR, and DE.NKR variants) were transiently expressed in HeLa cells, and GFP signal was monitored by confocal fluorescence microscopy. GFP signal (green channel) is shown in the ethanol-treated control condition and upon treatment with leptomycin B (LMB) (50 nM, 4.5 h). The scale bars in the pictures in the upper row correspond to 20 µm. The lower row shows a two-fold magnification of the indicated fields.

### The UPF1-NKR mutant exerts a dominant-negative eHect

While performing the experiments described above, we observed that the overexpression of the UPF1-NKR mutant was toxic to the cells. This observation, together with the fact that one of the three residues we mutated was shown to confer a dominant negative phenotype when mutated to cysteine [R843, (Sun et al. 1998)], prompted us to investigate if the UPF1-NKR mutant has a dominant negative effect. To test this, we transiently transfected HEK293 Flp-In T-Rex cells with plasmids encoding UPF1-GFP fusions, harboring either UPF1 WT or the DE, NKR, and DE.NKR mutant sequences, as well as GFP alone as a control. 48 h after transfection, NMD activity was assessed by measuring the levels of some endogenous NMD-sensitive transcripts by RT-qPCR, and the expression level of the overexpressed proteins was determined by western blotting (Fig. 5A-B). Consistent with our hypothesis, we observed that even in the presence of endogenous UPF1, the overexpression of the UPF1-NKR-GFP protein resulted in NMD inhibition, as evidenced by the specific accumulation of NMD-sensitive transcripts in this condition compared to the GFP overexpression control (Fig. 5A). Also in this transient system, overexpression of the UPF1-WT-GFP protein did not result in increased NMD activity, further strengthening the notion that UPF1 is not the rate-limiting NMD factor under these experimental conditions. However, the results obtained with the UPF1-DE and UPF1-DE.NKR mutants were unexpected, as neither of them inhibited NMD when transiently overexpressed in HEK cells (Fig. 5A-B).

**Figure 5.**
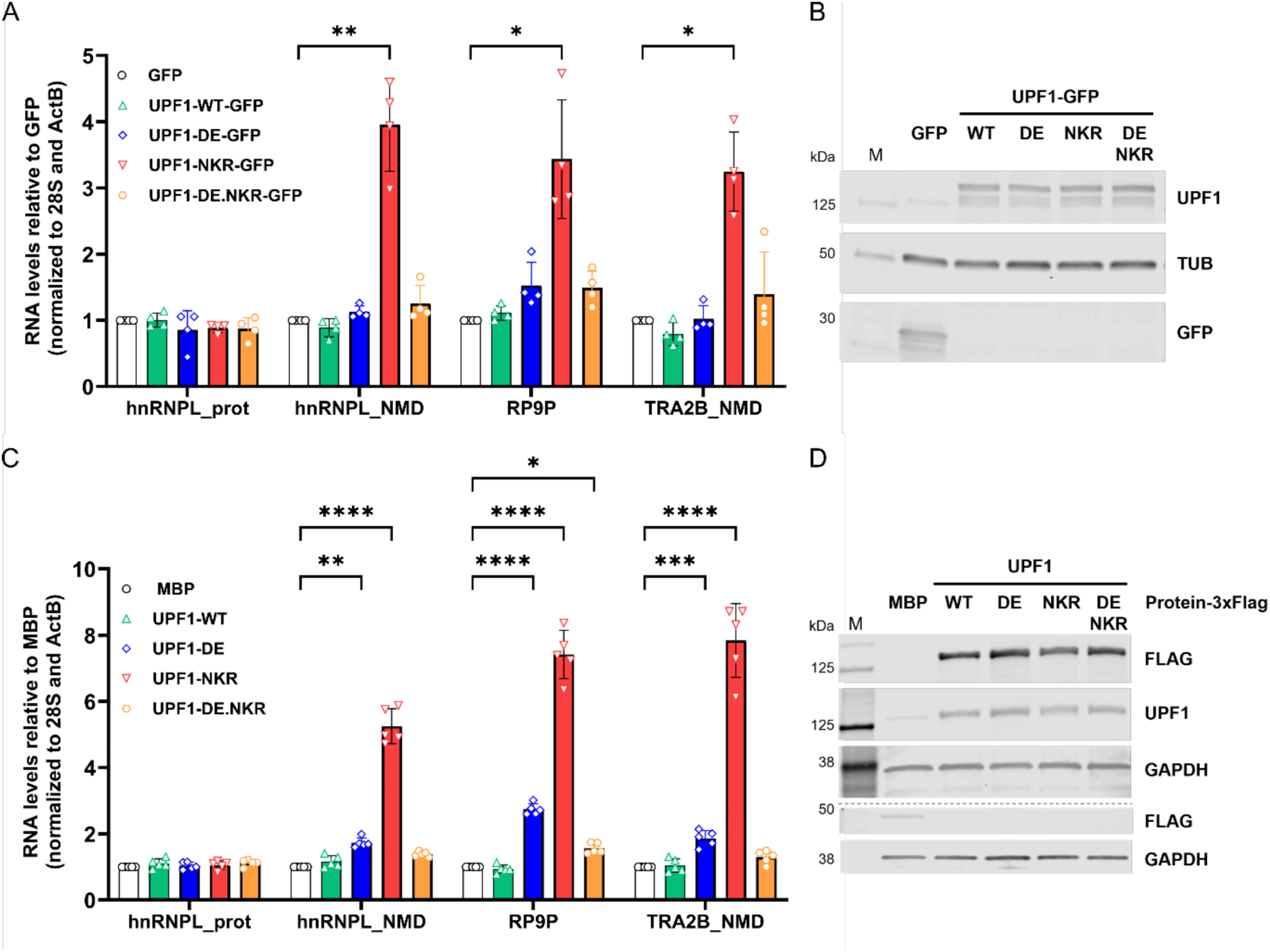
The UPF1-NKR mutant exerts a dominant-negative eHect. (A-B) HEK cells were transiently transfected to express UPF1-GFP (WT, DE, NKR, and DE.NKR variants) or GFP as a control. (A) Relative mRNA levels of hnRNPL_Protein, hnRNPL_NMD, RP9P, and TRA2B_NMD, normalized to 28S rRNA and actin mRNA levels, were determined by RT-qPCR 48 h after transfection. Averages and standard deviations from four independent experiments are shown. (B) The expression level of the overexpressed recombinant proteins was assessed by western blotting using the indicated antibodies. Tubulin served as loading control. M = molecular weight marker. (C-D) HEK cells stably transfected with pKK plasmids were induced for 48 h with Doxycycline to express UPF1-3xFlag proteins (WT, DE, NKR, or DE.NKR) or MBP-3xFlag as a control. (C) Relative mRNA levels were determined as in (A). Averages and standard deviations from five independent experiments are shown. (D) Western blotting assessing the expression of the overexpressed recombinant proteins as in (B), using the indicated antibodies. GAPDH served as loading control. Statistical significance in RT-qPCR data was determined by Two-way ANOVA and Dunnett’s multiple comparisons test against the GFP (A) or MBP (C) controls. Asterisks are only displayed for the comparisons with a *p*-value smaller than 0.05. * p ≤ 0.05; ** p ≤ 0.01; *** p ≤ 0.001; **** p ≤ 0.0001.

As an alternative method to verify these results, we stably integrated pKK plasmids into HEK293 Flp-In T-Rex cells to express a luciferase-targeting RNAi (CTRL KD) and C-terminally 3xFlag-tagged MBP, UPF1-WT, DE, NKR, or DE.NKR proteins, in a Dox-inducible manner. The expression of the recombinant proteins was induced for 48 h, and NMD activity was assessed by RT-qPCR. Also in this stable expression system, the UPF1-NKR mutant showed a strong dominant-negative effect (Fig. 5C-D). Contrary to the transient overexpression experiments (Fig. 5A), the stable expression of the UPF1-DE mutant resulted in a small but reproducible NMD inhibition (Fig. 5C-D). We attribute this difference to the more uniform expression of the recombinant proteins in the stable system, compared to the inherently more variable transient transfections. Strikingly, and consistent with the transient overexpression experiments, we did not observe NMD inhibition after the stable expression of the UPF1-DE.NKR double mutant (Fig. 5C).

Next, we set out to investigate the mechanism by which the UPF1-NKR mutant exerts its dominant negative effect on NMD. We speculated that the differential interaction of the UPF1 mutant proteins with endogenous NMD factors might explain their different dominant negative activities (or the lack thereof). We therefore tested the ability of the different UPF1 mutant proteins to interact with the NMD factors UPF2, UPF3B, and SMG6 by co-immunoprecipitation. To this end, we transiently overexpressed UPF1-WT-GFP, UPF1-DE-GFP, UPF1-NKR-GFP, and UPF1-DE.NKR-GFP proteins in HEK293 Flp-In T-Rex cells, followed by immunoprecipitation with anti-GFP antibodies and western blotting. We observed that all the UPF1-GFP overexpressed proteins co-immunoprecipitated UPF2, UPF3B, and SMG6 proteins (Suppl. Fig. 4A-B). The UPF1-DE mutant showed the strongest interaction with the tested NMD factors, consistent with previous reports showing that this mutant is trapped in NMD intermediate complexes (Franks et al. 2010). We found that UPF2, UPF3B, and SMG6 were pulled down with a similar efficiency by the UPF1-NKR and the UPF1-DE.NKR mutant proteins, despite their different NMD-inhibiting potential.

One key step for NMD activation is the phosphorylation of UPF1 (Kurosaki et al. 2014; Lee et al. 2015). We therefore tested whether stable expression of UPF1-3xFlag mutant proteins for 48 h resulted in altered UPF1 phosphorylation levels. Under our experimental conditions, we were unable to detect endogenous levels of phospho-UPF1; however, from the overexpressed proteins, we observed increased levels of phospho-UPF1 in the UPF1-DE overexpression condition (Suppl. Fig. 4C). Our results are consistent with the reported accumulation of the phosphorylated form of this mutant in stalled NMD intermediate complexes (Durand et al. 2016), and exclude changes in UPF1 phosphorylation as an explanation for the dominant-negative activity of UPF1-NKR. Finally, given the translation-dependent nature of NMD, we tested whether the overexpression of the UPF1 mutant proteins impaired cellular translation. To this end, we performed puromycin incorporation assays in HEK cells stably expressing 3xFlag-tagged UPF1 WT or mutant proteins. We did not observe any differences in the overall translation activities of the cells after 48 h of Dox-induced protein expression (Suppl. Fig. 4D), suggesting that translation inhibition cannot account for the dominant negative activity of the UPF1-NKR protein.

## Discussion

We describe here the design and characterization of a new RNA binding mutant of UPF1, termed UPF1-NKR, that harbors three point mutations, N524A, K547A, R843A. We show that these mutations affect UPF1’s capacity to interact with RNA *in vitro* and, consequently, also abrogate UPF1’s ATPase activity. Additionally, we demonstrate that UPF1-NKR is not only unable to restore NMD activity in UPF1-depleted cells but that it actually exerts a dominant negative effect on endogenous UPF1 when overexpressed. Finally, by studying the subcellular localization of various UPF1 mutants, we could determine that UPF1’s nuclear-cytoplasmic shuttling occurs independently of its ATPase or RNA-binding activities.

Our biochemical characterization of full-length hUPF1 purified from mammalian cells is a valuable addition to the large body of literature describing UPF1’s biochemical activities. Many studies have described UPF1’s interaction with nucleic acids in detail. The majority of these have characterized the binding of truncated versions of the protein, typically the helicase domain or the helicase domain together with the UPF2-interacting CH domain, purified from bacterial cells (Cheng et al. 2007; Chakrabarti et al. 2011; Fiorini et al. 2018; Gowravaram et al. 2018; Chapman et al. 2022). In addition, the nucleic acid binding capacity of full-length human UPF1 purified from insect cells (Bhattacharya et al. 2000) or from *E. coli* (Dehghani-Tafti and Sanders 2017) has also been reported. Several methods have been used in these reports to study the capacity of UPF1 to interact with DNA or RNA, including electrophoretic mobility shift assays (EMSA) (Bhattacharya et al. 2000; Dehghani-Tafti and Sanders 2017), fluorescence anisotropy (Chakrabarti et al. 2011; Gowravaram et al. 2018; Chapman et al. 2022), or label-free biosensing techniques (Cheng et al. 2007; Fiorini et al. 2018). The reported dissociation constants (*K_d_*) for UPF1 binding to nucleic acids range from ∼ 1-50 nM. This wide range of measured *K_d_* can be attributed to the wide spectrum of experimental conditions in which UPF1’s binding to nucleic acids has been tested. Nevertheless, a general conclusion can be made from these reports: UPF1 has a high affinity for single-stranded nucleic acids, with a *K_d_* in the lower nanomolar range, and its binding to nucleic acids is decreased in the presence of ATP. Our calculated dissociation constants for UPF1 binding to RNA of ∼16 nM and ∼873 nM in the absence and presence of ATP, respectively, corroborate this conclusion. Since we carefully tested the influence of the incubation time on the *K_d_* measurements, we are confident that our reported *K_d_* were measured at binding equilibrium (Jarmoskaite et al. 2020). On the other hand, we acknowledge that our *K_d_* values might be slightly overestimated since the sensitivity of our MST experiment precluded us from using RNA concentrations lower than 4 nM. However, differences between our UPF1 *K_d_* values and those previously reported, particularly those determined for truncated UPF1, might also stem from the presence of the N-and C-terminal domains, which were shown to exert auto-inhibitory functions on the helicase domain of UPF1 (Chamieh et al. 2008; Clerici et al. 2009; Chakrabarti et al. 2011; Fiorini et al. 2013).

We demonstrate that hUPF1 purified from mammalian cells can hydrolyze ATP *in vitro*, confirming, along with RNA binding experiments, that our purification protocol yields active UPF1 molecules. Our purification strategy begins with high salt (500 mM NaCl) and gradually lowers the concentration during subsequent washes, reaching a final, more physiological concentration of 150 mM NaCl. The initial high stringency, combined with a smooth transition to lower salt conditions, is crucial for obtaining pure, active UPF1. Purifying recombinant proteins from cells to test their activities *in vitro* bears the inherent potential risk of assaying the activities of co-purified proteins, especially when the target purified protein is an RNA-binding protein. We would therefore like to emphasize the importance of having control proteins that are purified from the same cells following the same protocol. In our case, we not only purified the unrelated protein MBP but also the ATPase-deficient UPF1-DE. This mutant UPF1 proved very useful since it retains RNA binding capacity but fails to hydrolyze ATP, features that were previously described, and gave us confidence that the measured activities were intrinsic to the purified UPF1 species.

The UPF1-NKR RNA binding mutant presented here showed no trace of RNA binding in the MST experiments and, correspondingly, also showed no ATPase activity *in vitro*. Notably, mutations in two of the three amino acids we altered were previously reported to affect UPF1’s binding to nucleic acids. Exchanging the lysine in position 547 (in human UPF1 isoform 2, K485) to a proline has been shown to reduce UPF1’s binding to nucleic acids *in vitro* in experiments using the helicase domain of both human (Chapman et al. 2022) or yeast UPF1 (Kanaan et al. 2018). Likewise, the point mutant R843C (yeast R779C) shows reduced nucleic acid binding and exerts a dominant-negative effect on endogenous UPF1 (Leeds et al. 1992; Sun et al. 1998; Kurosaki et al. 2014; Chapman et al. 2022). Each of these two mutations weakens UPF1 binding to nucleic acids, but, to our knowledge, they have previously never been tested in combination. The crystal structure of yeast Upf1 complexed to RNA and ADP:AlF_4_^-^ shows that UPF1 interacts with ∼8 nucleotides, with the arginine residue (yeast R779 / human R843) in the RecA2 domain contacting ribonucleotides towards the 5‘ end and the lysine residue in the RecA1 domain (yeast K485 / human K547) contacting ribonucleotides located in the 3’ end of the RNA (Chakrabarti et al. 2011). In addition to these two residues, the NKR mutant also harbors a modification in residue N524, which contacts the central nucleotides in the RNA molecule (yeast N462 in the structure). The combined destabilization of the UPF1-RNA interaction at three different positions along the binding surface results in the severe reduction in RNA binding of UPF1-NKR.

In this work, we characterized the UPF1-NKR mutant as a strong dominant-negative protein, capable of inhibiting NMD when overexpressed in mammalian cells. The extent of NMD inhibition achieved by stable expression of the UPF1-NKR mutant for 48 h is greater than that observed after inducing an shRNA-mediated knockdown of endogenous UPF1 for the same time (Fig. 5C and Suppl. Fig. 2). This makes UPF1-NKR overexpression an interesting and easy orthogonal method to inhibit NMD, similar to UPF1 R843C (Sun et al. 1998). Along with the UPF1-NKR, we also tested the capacity of the other two mutants to inhibit NMD. We predicted that the UPF1-DE mutant would also act as a dominant negative, as its failure to hydrolyze ATP renders it unable to dissociate from RNAs and leads to its accumulation in P bodies as part of stalled intermediate decay complexes (Franks et al. 2010). However, we found that the stable overexpression of the UPF1-DE mutant only mildly inhibits NMD. We speculate that if the UPF1-DE can occasionally dissociate from RNA, mRNA decay could proceed. This is consistent with a transcriptomic analysis showing that the loss of NMD activity due to a UPF1 knockdown can be partially rescued by expression of the UPF1-DE mutant (Chapman et al. 2024).

Even more surprising were the results obtained with the UPF1-DE.NKR double mutant, as the addition of the DE mutation to the dominant dominant-negative UPF1-NKR mutation abolished the dominant-negative capacity conferred by the NKR mutation. A straightforward hypothesis is that the UPF1-NKR could titrate away NMD factors, rendering them limiting to interact with endogenous UPF1. We tested this hypothesis by co-immunoprecipitation experiments. Among the three UPF1 mutants, UPF1-DE showed the strongest interaction with UPF2, UPF3B, or SMG6, while UPF1-NKR and UPF1-DE.NKR co-immunoprecipitated less of these NMD factors, arguing against a simple sequestering mechanism. However, it is plausible that the NMD factors might have to interact with their partner proteins in a specific spatial arrangement and in the correct temporal order, and the co-immunoprecipitations might fail to capture such putative differences between the UPF1 mutants. The interactions of the endogenous NMD factors with the UPF1 mutants could have distinct functional consequences: Those involving the UPF1-DE mutant are likely to occur while bound to mRNA, whereas those involving the UPF1-NKR mutant would take place in the absence of RNA. Yet, the conundrum of the UPF1-DE.NKR double mutant remains. We confirmed that UPF1-NKR and UPF1-DE.NKR cannot bind RNA *in vitro*. However, it would be interesting to know whether this is also the case in a cellular context and for specific transcripts. It is conceivable that compared to UPF1-NKR, the UPF1-DE.NKR mutant may still weakly bind RNA in a physiological context, either assisted by other cellular RNA-binding proteins or facilitated by the DE mutation, which by itself causes UPF1 to remain bound to RNA. Even though our results do not explain the mechanism underlying the dominant-negative nature of the UPF1-NKR, our biochemical characterization of UPF1-DE and UPF1-DE.NKR highlights the role of tight RNA binding of UPF1-DE for its entrapment in P bodies, its stronger interaction with other NMD factors, and its accumulation in a hyperphosphorylated form (Franks et al. 2010).

UPF1 shuttling (Mendell et al. 2002) and the reported cytoplasmic and nuclear functions that depend on its biochemical activities [reviewed in (Azzalin and Lingner 2006a; Imamachi et al. 2012)] suggested that UPF1 shuttles between the nucleus and cytoplasm in a manner regulated by, or dependent on, these activities. This hypothesis was tested in *Drosophila* S2 cells and larvae (Singh et al. 2019). Immunofluorescence and fractionation experiments showed that at steady state, only a small proportion of UPF1 localized to the nucleus and that blocking CRM1-dependent nuclear export by LMB treatment resulted in a rapid accumulation of UPF1 in the nucleus. To test if UPF1’s helicase activity was required for its shuttling, the subcellular localization of the ATPase-deficient UPF1-DE617AA (DE636AA in human UPF1) fused to GFP was determined, and it was found that, contrary to WT UPF1, the UPF1-DE mutant was unable to localize to the nucleus after LMB treatment and stayed in bright cytoplasmic granules. These results were interpreted as evidence for UPF1’s helicase activity being required for its nuclear-cytoplasmic shuttling (Singh et al. 2019). Our results from experiments in mammalian cells expressing the UPF1-DE mutant are in complete agreement with those performed in *Drosophila* cells. However, the normal shuttling behavior exhibited by the UPF1-NKR mutant, which like UPF1-DE is unable to hydrolyze ATP, led us to revisit the notion that ATP hydrolysis was required for UPF1’s shuttling. We reasoned that since ATP hydrolysis is required for UPF1 to dissociate from nucleic acids, the tight binding to RNA exhibited by the UPF1-DE protein might anchor it to cytoplasmic P bodies, rendering it unable to shuttle to the nucleus. We confirmed this hypothesis by showing that the double mutant UPF1-DE.NKR, which failed to bind RNA *in vitro*, no longer localized to P bodies and regained normal nuclear-cytoplasmic shuttling. Overall, our results show that UPF1’s nuclear-cytoplasmic shuttling is not determined by its RNA binding or ATPase activities, and hence most likely also not by its helicase activity. Our results further indicate that only RNA-free UPF1 can shuttle to the nucleus. We speculate that at any given time, most cellular UPF1 might be bound to RNA in the cytoplasm of mammalian cells, considering UPF1’s high affinity for RNA and the recent estimates that UPF1 and mRNA molecules are present at about 1:1 stoichiometry (Cho et al. 2022; Chapman et al. 2024). Consequently, only a small proportion of RNA-free UPF1 would be available for shuttling to the nucleus. This small amount of shuttling UPF1, and the presence of a strong nuclear export signal (NES) in UPF1 (Eberle et al. 2021), might explain why only a very small, hardly detectable fraction of UPF1 can be found in the nucleus of mammalian cells at steady state. Our results are consistent with scenarios where UPF1’s trafficking to the nucleus is driven by intrinsic sequence or structural features of UPF1, or by direct protein interaction partners. Future studies aimed at dissecting the nuclear import pathway(s) of UPF1 will be key to finally tease apart the alleged nuclear functions of UPF1 from those related to UPF1’s involvement in cytoplasmic RNA decay pathways.

## Material and Methods

### Plasmids

The oligonucleotides used are listed in Table S1. To generate the UPF1-WT-GFP-expressing plasmid, the GFP coding sequence was inserted, replacing the FLAG coding sequence within the pcDNA3-NG-UPF1-WT-FLAG plasmid (Rufener and Muhlemann 2013) using *Not* I and *Xba* I restriction enzymes. Annealed oligonucleotides encoding a linker of 16 glycine residues (see sequence in Table S1) were inserted into the vector at the *Not* I site to generate the plasmid encoding the fusion protein UPF1-GFP with a G16 flexible linker in between. UPF1-DE-GFP was generated by exchanging a fragment of UPF1-WT-GFP (from *Kpn* I to *Sbf* I) with the equivalent from pcDNA3-NG-UPF1-WT-FLAG DE636AA containing the two desired changes. Point mutations for the UPF1-NKR-GFP were introduced individually by site-directed mutagenesis PCR on cloning plasmids containing fragments of UPF1. The mutated UPF1 sequences were then cloned into the final plasmid by restriction sites (oligonucleotides for point mutations are listed in Table S1). To construct the UPF1-DE.NKR-GFP mutant, a fragment from pcDNA3-NG-UPF1-WT-FLAG DE636AA containing the two desired changes was subcloned into the UPF1-NKR-GFP plasmid, using *Kpn* I and *Sbf* I restriction enzymes.

pKK plasmids expressing from bidirectional, Tet responsive CMV promoter an RNAi cassette (targeting UPF1 or luciferase as control) on one side and the protein of interest (triple FLAG-tagged MBP or UPF1) on the other side were designed and cloned based on the original plasmid (Szczesny et al. 2018). Each RNAi cassette consists of three siRNAs, which contain self-complementary regions and a loop-forming sequence from murine miRNA 155. For designing the RNAi cassette against (firefly) luciferase (*P. pyralis*, NCBI accession number M15077.1) LOCK-iT™ RNAi Designer from Thermo Fisher Scientific (https://bit.ly/3qcESry) was used, and for the UPF1 RNAi cassette, previously established siRNA target sequences were chosen (Paillusson et al. 2005). The RNAi cassettes were synthesized (General Biosystems, Inc.) and inserted into the pKK vectors using *Apa* I and *Mfe* I restriction sites. C-terminally triple FLAG-tagged MBP was inserted into pKK plasmids using InFusion cloning (Takara). To introduce the WT UPF1 sequence with a C-terminal 3xFlag tag, the pKK vector was partially digested with *Not* I and *Hind* III (since two *Not* I sites are present), and the desired vector backbone was purified via agarose gel. RNAi-resistant WT UPF1 coding sequence was obtained from the pcDNA3-NG-UPF1-WT-FLAG (Rufener and Muhlemann 2013) plasmid using *Not* I and *Hind* III and cloned into the linearized vector backbone. To obtain pKK plasmids with variant UPF1 (DE, NKR, and DE.NKR), the corresponding fragments of the WT UPF1 sequence were replaced by the mutant ones using *Hind* III and *Pml* I restriction sites.

For all PCR reactions, high-fidelity polymerases were used (Kapa Hifi Hot Start Ready mix (Roche) or CloneAmp HiFi PCR Premix (Takara)) and the sequences of all plasmids were confirmed by Sanger sequencing (Microsynth, Switzerland).

### Cell culture and transfection

HeLa cells and HEK293 Flp-In^TM^ T-Rex cells (Thermo Fisher Scientific) were cultured in Dulbecco’s Modified Eagle Medium (DMEM) supplemented with 10% fetal bovine serum (FBS), 100 U/ml penicillin and 100 μg/ml streptomycin (P/S) (DMEM^+/+^) at 37 °C under 5% CO_2_.

To stably introduce pKK plasmids into HEK293 Flp-In^TM^ T-Rex cells, 3 x 10^6^ cells were seeded in a 10 cm plate using DMEM plus tetracycline-free FBS. The next day, 2 μg of the pKK plasmid and 18 μg of pOG44 Flp-Recombinase expression vector (Thermo Fisher Scientific) were transfected using Lipofectamine 2000 (Thermo Fisher Scientific). 24 h after transfection, cells were transferred into T150 flasks (DMEM + tetracycline-free FBS), and 3 h later the antibiotics were added (P/S, 100 μg/ml hygromycin B, 10 μg/ml blasticidine). Cells were kept under selection for about two weeks. Transcription of the plasmids was induced by the addition of 1 μg/ml doxycycline (Dox), and cells were harvested at the indicated time points after Dox-induction.

For transient transfection in HEK293 Flp-In^TM^ T-Rex cells, 6 x 10^5^ cells per well were seeded in a 6-well plate and transfected the following day with 500 ng/well of GFP or UPF1-GFP expressing plasmids using Dogtor transfection reagent (OZ Biosciences). Cells were harvested 48 h after transfection.

HEK cells transiently transfected with UPF1-GFP expression plasmids, or stably transfected with pKK plasmids, were harvested for RNA analysis using a guanidinium thiocyanate-phenol-chloroform mix (TRI-mix, (Nicholson et al. 2012)) or the Maxwell® RSC simplyRNA Cells Kit (Promega), and for protein analysis by preparing cell lysates using RIPA buffer (50 mM Tris-HCl (pH 8.0), 1 mM EDTA, 1% Triton X-100, 0.25% sodium deoxycholate, 150 mM NaCl) supplemented with protease and phosphatase inhibitors.

For the puromycin incorporation assay, HEK293 Flp-In^TM^ T-Rex cells stably transfected with pKK plasmids harboring a luciferase-targeting RNAi cassette (CTRL KD) and MBP-3xFlag or UPF1-3xFlag (WT, DE, NKR, or DE.NKR) expression constructs were seeded into 6-well plates at a density of 2×10^5^ cells/well in DMEM^+/+^ supplemented with 1 μg/ml Doxycyclin. 48 h later, the medium was changed to DMEM^+/+^ with 10 μg/ml of puromycin for 10 min. Subsequently, the cells were washed twice with 1x PBS, and then allowed to recover in DMEM^+/+^ with 1 μg/ml Doxycyclin for 30 min. As a positive control for translation inhibition, one well of every cell line was treated with 100 μg/ml Cycloheximide for 2 h before the puromycin pulse, as well as during the 30-minute recovery after it. Samples were collected for protein analysis via western blotting using RIPA buffer supplemented with protease and phosphatase inhibitors.

For transient transfection in HeLa cells, 4 x 10^5^ cells per well were seeded in a 6-well plate and transfected the following day with 500 ng/well of GFP or UPF1-GFP expressing plasmids using Dogtor transfection reagent (OZ Biosciences). 24 h later, transfection efficiencies were evaluated by using fluorescence microscopy and 40’000 cells per well were seeded in 8-well chamber slides for fluorescence microscopy analysis the next day.

### LMB treatment and fluorescence microscopy analysis

Cells in 8-well chamber slides were incubated for 4.5 h with 50 nM LMB (Cell Signaling Technology) or ethanol as control, followed by two washes with PBS and fixation with 4 % paraformaldehyde for 25 minutes at room temperature. Next, cells were washed three times with TBS (20 mM Tris-HCl, pH 7.5, 150 mM NaCl) and incubated with TBS +/+ (TBS plus 0.5 % Triton X-100 and 6 % FBS) containing DAPI for 30 minutes at 37 °C. Finally, cells were washed three times with TBS, and Vectashield mounting medium (Vector Laboratories) was added on slides to mount the coverslips. The slides were analyzed by confocal spinning disc microscopy (Nikon TI2 with Crest X-Light V2).

### Co-immunoprecipitation

For investigating interaction partners of UPF1, 4 μg of UPF1-GFP (WT, DE, NKR, or DE.NKR) or GFP-expressing plasmid (as control) were transfected with Dogtor transfection reagent (OZ Biosciences) into HEK293 Flp-In^TM^ T-Rex cells seeded the previous day in a 15 cm plate. 48 h after transfection, cells were harvested and lysed in cold lysis buffer (50 mM HEPES, pH 7.3, 150 mM NaCl, 0.5 % Triton X-100 with protease and phosphatase inhibitors) using dual centrifugation for 4 minutes at 1500 rpm and-5 °C (Zentrimix 380R, Hettich). After centrifugation (16’000 x g, 10 min, 4° C), the supernatant was incubated with 25 μl of GFP-Trap magnetic agarose beads (Chromotek) on a rotating wheel for one hour at 4 °C. After washing the beads twice on a magnet with lysis buffer, the samples were treated with 100 μg RNase A for 10 minutes at 25 °C. The samples were washed once with lysis buffer and eluted by adding SDS-loading buffer to the beads. The samples were incubated for 10 minutes at 98 °C and analyzed via Western blotting.

### Western blot analysis

Protein samples were loaded into 4-15% Criterion™ TGX™ Precast gels (Bio-Rad) or mPAGE® 4-12% Bis-Tris precast gels (Millipore) and electrophoresed in 1x SDS-PAGE running buffer or 1x MOPS running buffer, respectively. After electrophoresis, proteins were transferred onto nitrocellulose membranes using the Trans-Blot Turbo Transfer Packs and either the High MW or the Mixed MW protocols preloaded in the Trans-Blot turbo transfer system (Bio-Rad). Efficient protein transfer to the membranes was verified using the Revert™ 700 total protein stain (LI-COR), following the manufacturer’s instructions.

Membranes were blocked for 1 h at room temperature (RT) in 5% BSA-TBS-T (0.1% Tween20 in TBS buffer) and then incubated with the indicated primary antibodies for 2 h at RT or overnight at 4°C. Primary antibodies were diluted in 5% BSA-TBS-T. After three 10-minute washes in TBS-T, membranes were incubated with fluorescently labeled secondary antibodies in 5% BSA-TBS-T for 1 h at RT, followed by three 10-minute TBS-T washes. The blots were then visualized using the Odyssey Imaging System (LI-COR).

Antibodies used in this study: Mouse anti-Flag (Sigma, F1804); Goat anti-UPF1(Bethyl Laboratories, A300-038A); Goat anti-GFP (Acris Antibodies, AB0020-200); Mouse anti-Tubulin (Sigma, T9028); Mouse anti-GAPDH (Santa Cruz Biotechnology, sc-47724); Rabbit anti-UPF2 (Bethyl Laboratories, A303-929A); Rabbit anti-UPF3B (Abcam, ab134566); Rabbit anti-SMG6 (Abcam, ab87539); Rabbit anti-phospho-UPF1 (Ser1127) (Merck Millipore, 07-1016); Mouse anti-puromycin (Millipore, MABE343).

### RNA analysis by RT-qPCR

Total RNA was extracted using TRI-mix and precipitated with isopropanol as described previously (Nicholson et al. 2012), or by using the Maxwell® RSC simplyRNA Cells Kit (Promega). RNA concentration was measured by Nanodrop, and 1 μg of RNA was reverse transcribed using random hexamers and the AffinityScript Multi-Temp Reverse Transcriptase (Agilent), following the manufacturer’s protocol. 15 µl qPCR reactions were set with 24 ng of cDNA and 500 nM of each primer, in 1xBrilliant III Ultra-Fast SYBR Green qPCR mix (Agilent). cDNA levels were measured with the Rotor-Gene Q (Qiagen), running technical duplicates for every sample. The absence of DNA contamination was controlled by running RT minus controls in each experiment. Relative quantitation was performed using the ΔΔCt method. Oligonucleotides for qPCRs are listed in Table S1.

### Purification of recombinant proteins

Triple Flag-tagged UPF1-WT, UPF1-DE, UPF1-NKR, UPF1-DE.NKR, and MBP proteins were expressed in HEK293 Flp-In^TM^ T-Rex cells stably transfected with the respective pKK plasmids and purified by immunoprecipitation. Expression of the recombinant proteins was induced by the addition of 1 µg/ml Dox for 48 h. Cell pellets containing each 6 x 10^7^ cells were resuspended in 700 μl lysis/wash buffer 1 (500 mM NaCl, 0.5 % Triton-X 100, 50 mM Hepes pH 7.4, supplemented with protease and phosphatase inhibitors) and lysed by dual centrifugation for 4 minutes at 1500 rpm and-5 °C (Zentrimix 380R, Hettich). The cell lysate was clarified by centrifugation (16’000 x g, 4 °C, 10 min), and the supernatant was incubated with 60 μl Dynabeads M-270 Epoxy beads (Invitrogen) coupled with anti-FLAG M2 antibodies (Sigma) for 1 h at 4 °C on a rotating wheel. The beads were washed once with 1 ml lysis/ wash buffer 1, once with 1 ml wash buffer 2 (250 mM NaCl, 0.25 % Triton-X 100, 25 mM Hepes pH 7.4, supplemented with protease and phosphatase inhibitors), and once with 1 ml wash buffer 3 (150 mM NaCl, 0.1 % Triton-X 100, 10 mM Hepes pH 7.4, supplemented with protease and phosphatase inhibitors). The proteins were eluted by adding 50 μl elution buffer (1 mg/ml 3xFLAG peptide, 150 mM NaCl, 0.1 % Triton-X 100, 10 mM Hepes pH 7.4, supplemented with protease and phosphatase inhibitors) and incubating it for 15 minutes at 25 °C at 1000 rpm on a thermomixer. An SDS-PAGE followed by Imperial Protein Stain (Thermo Scientific) staining controlled for the purity of the eluted proteins. To estimate the concentration of the purified proteins, a BSA standard curve was loaded on the same gel.

### MicroScale Thermophoresis

The RNA binding activity of the purified recombinant proteins was assessed using MicroScale Thermophoresis (MST). The 5‘-labeled Cy5 RNA U30 oligonucleotide (Microsynth) was diluted to 8 nM in 1x elution buffer (150 mM NaCl, 0.1 % Triton-X 100, 10 mM Hepes pH 7.4, supplemented with protease and phosphatase inhibitors). The unlabeled recombinant proteins were serially diluted from a concentration of 300 nM down to 0.009 nM in 1x elution buffer (150 mM NaCl, 0.1 % Triton-X 100, 10 mM Hepes pH 7.4, supplemented with protease and phosphatase inhibitors) and mixed with a constant amount of labeled RNA (final concentration of 4 nM). Binding reactions were incubated at 25 °C for the indicated time periods in the presence or absence of 1 mM ATP. The RNA-protein mixtures were analyzed in standard capillaries using the Monolith NT.115T MST (NanoTemper Technologies). Measurements were carried out at 25 °C, with the following settings: LED power 100%, MST power 40%, laser on time 30 sec, laser oa time 5 sec. The NanoTemper Technologies Analysis software, MO.Affinity Analysis v2.3, was used to determine the corresponding *K_d_* using the *K_d_* fit model.

### NADH-coupled ATPase assay

The capacity of the purified proteins to hydrolyze ATP *in vitro* was tested using NADH-coupled ATPase assays, following a protocol previously reported (Fritz et al. 2020). Briefly, 60 μl reactions were assembled in a step-wise manner in a clear flat-bottomed 96-well plate. In the first step, a mix of 80 nM of the recombinant protein, 1x ATPase buffer (50 mM MES pH 6.0, 50 mM KOAc, 0.1 mM EDTA), and 0.4 mg/ml Poly(U) RNA was made, filled up to 30 μl with nuclease-free water, and incubated for 10 min at room temperature. Next, 24 μl of a reaction mix (1.25 x ATPase buffer, 5 mM MgOAc, 1.25 mM PEP, 1 mM NADH, 24.1 U/ml pyruvate kinase, 16.2 U/ml lactate dehydrogenase, filled up with nuclease-free water) was added to each well. Finally, to start the reaction, 6 μl 10 mM ATP was added and mixed by pipetting, and the reactions were incubated at 37 °C for 120 min in a Tecan M1000 plate reader, where the oxidation of NADH was followed by measuring the absorbance at 340 nm every minute.

### Graphs and statistical analysis

The figure with UPF1’s structure was generated using PyMol. Graphs displaying MST traces were generated using the NanoTemper Technologies software, MO.Affinity Analysis v2.3. All other graphs and statistical analyses were done with GraphPad Prism 10 software.

## Acknowledgments

We are grateful to Julia Schnider and Lena Grollmus for the UPF1-GFP and pKK-miRLuc plasmids, respectively. We also thank Dr. Hervé Le Hir for supplying the UPF1-HD purified protein, Dr. Christoph von Ballmoos and Dr. Lukas Rimle for assistance with the ATPase assays, and Nicole Kleinschmidt for her excellent technical support.

---

**Table S1.**
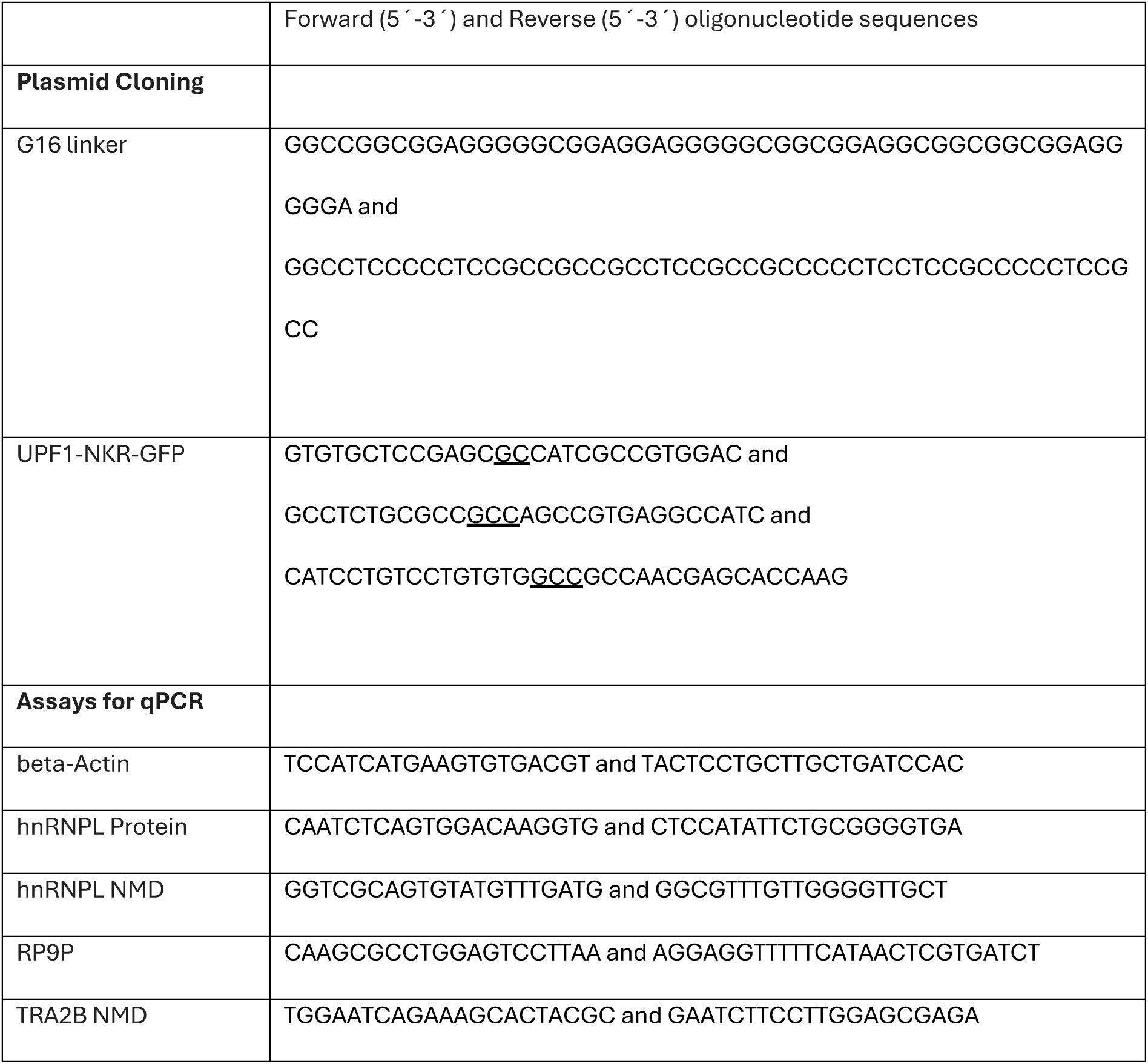
Oligonucleotides (from Microsynth, Switzerland) used in the study. Changes to alanines are underlined.

## References

Ajamian L, Abel K, Rao S, Vyboh K, Garcia-de-Gracia F, Soto-Rifo R, Kulozik AE, Gehring NH, Mouland AJ. 2015. HIV-1 Recruits UPF1 but Excludes UPF2 to Promote Nucleocytoplasmic Export of the Genomic RNA. Biomolecules 5: 2808–2839.

Applequist SE, Selg M, Raman C, Jack HM. 1997. Cloning and characterization of HUPF1, a human homolog of the Saccharomyces cerevisiae nonsense mRNA-reducing UPF1 protein. Nucleic Acids Res 25: 814--821.

Atkin AL, Altamura N, Leeds P, Culbertson MR. 1995. The majority of yeast UPF1 co-localizes with polyribosomes in the cytoplasm. Mol Biol Cell 6: 611–625.

Azzalin CM, Lingner J. 2006a. The double life of UPF1 in RNA and DNA stability pathways. Cell Cycle 5: 1496–1498.

Azzalin CM, Lingner J. 2006b. The human RNA surveillance factor UPF1 is required for S phase progression and genome stability. Curr Biol 16: 433–439.

Azzalin CM, Reichenbach P, Khoriauli L, Giulotto E, Lingner J. 2007. Telomeric repeat containing RNA and RNA surveillance factors at mammalian chromosome ends. Science 318: 798–801.

Bhattacharya A, Czaplinski K, Trifillis P, He F, Jacobson A, Peltz SW. 2000. Characterization of the biochemical properties of the human Upf1 gene product that is involved in nonsense-mediated mRNA decay. RNA 6: 1226–1235.

Carastro LM, Tan CK, Selg M, Jack HM, So AG, Downey KM. 2002. Identification of delta helicase as the bovine homolog of HUPF1: demonstration of an interaction with the third subunit of DNA polymerase delta. Nucleic Acids Res 30: 2232–2243.

Chakrabarti S, Jayachandran U, Bonneau F, Fiorini F, Basquin C, Domcke S, Le Hir H, Conti E. 2011. Molecular mechanisms for the RNA-dependent ATPase activity of Upf1 and its regulation by Upf2. Mol Cell 41: 693–703.

Chamieh H, Ballut L, Bonneau F, Le Hir H. 2008. NMD factors UPF2 and UPF3 bridge UPF1 to the exon junction complex and stimulate its RNA helicase activity. Nat Struct Mol Biol 15: 85–93.

Chapman JH, Craig JM, Wang CD, Gundlach JH, Neuman KC, Hogg JR. 2022. UPF1 mutants with intact ATPase but deficient helicase activities promote efficient nonsense-mediated mRNA decay. Nucleic Acids Res.

Chapman JH, Youle AM, Grimme AL, Neuman KC, Hogg JR. 2024. UPF1 ATPase autoinhibition and activation modulate RNA binding kinetics and NMD efficiency. Nucleic Acids Res 52: 5376–5391.

Chawla R, Redon S, Raftopoulou C, Wischnewski H, Gagos S, Azzalin CM. 2011. Human UPF1 interacts with TPP1 and telomerase and sustains telomere leading-strand replication. EMBO J 30: 4047–4058.

Cheng Z, Muhlrad D, Lim MK, Parker R, Song H. 2007. Structural and functional insights into the human Upf1 helicase core. EMBO J 26: 253–264.

Cho NH, Cheveralls KC, Brunner AD, Kim K, Michaelis AC, Raghavan P, Kobayashi H, Savy L, Li JY, Canaj H et al. 2022. OpenCell: Endogenous tagging for the cartography of human cellular organization. Science 375: eabi6983.

Clerici M, Mourao A, Gutsche I, Gehring NH, Hentze MW, Kulozik A, Kadlec J, Sattler M, Cusack S. 2009. Unusual bipartite mode of interaction between the nonsense-mediated decay factors, UPF1 and UPF2. EMBO J 28: 2293–2306.

Czaplinski K, Weng Y, Hagan KW, Peltz SW. 1995. Purification and characterization of the Upf1 protein: a factor involved in translation and mRNA degradation. RNA 1: 610–623.

Dehghani-Tafti S, Sanders CM. 2017. DNA substrate recognition and processing by the full-length human UPF1 helicase. Nucleic Acids Res 45: 7354–7366.

Durand S, Franks TM, Lykke-Andersen J. 2016. Hyperphosphorylation amplifies UPF1 activity to resolve stalls in nonsense-mediated mRNA decay. Nature communications 7: 12434.

Eberle AB, Lykke-Andersen S, Muhlemann O, Jensen TH. 2009. SMG6 promotes endonucleolytic cleavage of nonsense mRNA in human cells. Nat Struct Mol Biol 16: 49–55.

Eberle AB, Schranz K, Nasif S, Grollmus L, Mühlemann O. 2021. Dissecting the import and export pathways of the human RNA helicase UPF1. bioRxiv preprint.

Fiorini F, Bagchi D, Le Hir H, Croquette V. 2015. Human Upf1 is a highly processive RNA helicase and translocase with RNP remodelling activities. Nature communications 6: 7581.

Fiorini F, Boudvillain M, Le Hir H. 2013. Tight intramolecular regulation of the human Upf1 helicase by its N-and C-terminal domains. Nucleic Acids Res 41: 2404–2415.

Fiorini F, Robin JP, Kanaan J, Borowiak M, Croquette V, Le Hir H, Jalinot P, Mocquet V. 2018. HTLV-1 Tax plugs and freezes UPF1 helicase leading to nonsense-mediated mRNA decay inhibition. Nature communications 9: 431.

Franks TM, Singh G, Lykke-Andersen J. 2010. Upf1 ATPase-dependent mRNP disassembly is required for completion of nonsense-mediated mRNA decay. Cell 143: 938–950.

Fritz SE, Ranganathan S, Wang CD, Hogg JR. 2020. The RNA-binding protein PTBP1 promotes ATPase-dependent dissociation of the RNA helicase UPF1 to protect transcripts from nonsense-mediated mRNA decay. J Biol Chem 295: 11613–11625.

Gatfield D, Unterholzner L, Ciccarelli FD, Bork P, Izaurralde E. 2003. Nonsense-mediated mRNA decay in Drosophila: at the intersection of the yeast and mammalian pathways. EMBO J 22: 3960–3970.

Gowravaram M, Bonneau F, Kanaan J, Maciej VD, Fiorini F, Raj S, Croquette V, Le Hir H, Chakrabarti S. 2018. A conserved structural element in the RNA helicase UPF1 regulates its catalytic activity in an isoform-specific manner. Nucleic Acids Res 46: 2648–2659.

Huang L, Lou CH, Chan W, Shum EY, Shao A, Stone E, Karam R, Song HW, Wilkinson MF. 2011. RNA Homeostasis Governed by Cell Type-Specific and Branched Feedback Loops Acting on NMD. Mol Cell 43: 950–961.

Huntzinger E, Kashima I, Fauser M, Sauliere J, Izaurralde E. 2008. SMG6 is the catalytic endonuclease that cleaves mRNAs containing nonsense codons in metazoan. RNA 14: 2609–2617.

Imamachi N, Tani H, Akimitsu N. 2012. Up-frameshift protein 1 (UPF1): multitalented entertainer in RNA decay. Drug Discov Ther 6: 55–61.

Jarmoskaite I, AlSadhan I, Vaidyanathan PP, Herschlag D. 2020. How to measure and evaluate binding affinities. eLife 9.

Kanaan J, Raj S, Decourty L, Saveanu C, Croquette V, Le Hir H. 2018. UPF1-like helicase grip on nucleic acids dictates processivity. Nature communications 9: 3752.

Karousis ED, Muhlemann O. 2022. The broader sense of nonsense. Trends Biochem Sci 47: 921–935.

Kim YK, Maquat LE. 2019. UPFront and center in RNA decay: UPF1 in nonsense-mediated mRNA decay and beyond. RNA.

Kishor A, Fritz SE, Hogg JR. 2019. Nonsense-mediated mRNA decay: The challenge of telling right from wrong in a complex transcriptome. Wiley Interdiscip Rev RNA 10: e1548.

Kudo N, Wolff B, Sekimoto T, Schreiner EP, Yoneda Y, Yanagida M, Horinouchi S, Yoshida M. 1998. Leptomycin B inhibition of signal-mediated nuclear export by direct binding to CRM1. Exp Cell Res 242: 540–547.

Kurosaki T, Li W, Hoque M, Popp MW, Ermolenko DN, Tian B, Maquat LE. 2014. A post-translational regulatory switch on UPF1 controls targeted mRNA degradation. Genes Dev 28: 1900–1916.

Lee SR, Pratt GA, Martinez FJ, Yeo GW, Lykke-Andersen J. 2015. Target Discrimination in Nonsense-Mediated mRNA Decay Requires Upf1 ATPase Activity. Mol Cell 59: 413–425.

Leeds P, Peltz SW, Jacobson A, Culbertson MR. 1991. The product of the yeast UPF1 gene is required for rapid turnover of mRNAs containing a premature translational termination codon. Genes Dev 5: 2303–2314.

Leeds P, Wood JM, Lee BS, Culbertson MR. 1992. Gene products that promote mRNA turnover in Saccharomyces cerevisiae. Mol Cell Biol 12: 2165–2177.

Lloyd JPB. 2018. The evolution and diversity of the nonsense-mediated mRNA decay pathway. F1000Research 7: 1299.

Longman D, Jackson-Jones KA, Maslon MM, Murphy LC, Young RS, Stoddart JJ, Hug N, Taylor MS, Papadopoulos DK, Caceres JF. 2020. Identification of a localized nonsense-mediated decay pathway at the endoplasmic reticulum. Genes Dev 34: 1075–1088.

Lykke-Andersen J, Shu MD, Steitz JA. 2000. Human Upf proteins target an mRNA for nonsense-mediated decay when bound downstream of a termination codon. Cell 103: 1121–1131.

Mendell JT, ap Rhys CM, Dietz HC. 2002. Separable roles for rent1/hUpf1 in altered splicing and decay of nonsense transcripts. Science 298: 419–422.

Nasif S, Contu L, Muhlemann O. 2017. Beyond quality control: The role of nonsense-mediated mRNA decay (NMD) in regulating gene expression. Semin Cell Dev Biol.

Nicholson P, Joncourt R, Muhlemann O. 2012. Analysis of nonsense-mediated mRNA decay in mammalian cells. Curr Protoc Cell Biol Chapter 27: Unit27 24.

Okada-Katsuhata Y, Yamashita A, Kutsuzawa K, Izumi N, Hirahara F, Ohno S. 2012. N-and C-terminal Upf1 phosphorylations create binding platforms for SMG-6 and SMG-5:SMG-7 during NMD. Nucleic Acids Res 40: 1251–1266.

Paillusson A, Hirschi N, Vallan C, Azzalin CM, Muhlemann O. 2005. A GFP-based reporter system to monitor nonsense-mediated mRNA decay. Nucleic Acids Res 33: e54.

Perlick HA, Medghalchi SM, Spencer FA, Kendzior RJ, Jr., Dietz HC. 1996. Mammalian orthologues of a yeast regulator of nonsense transcript stability. Proc Natl Acad Sci U S A 93: 10928–10932.

Pulak R, Anderson P. 1993. mRNA surveillance by the Caenorhabditis elegans smg genes. Genes Dev 7: 1885–1897.

Rufener SC, Muhlemann O. 2013. eIF4E-bound mRNPs are substrates for nonsense-mediated mRNA decay in mammalian cells. Nat Struct Mol Biol 20: 710–717.

Sato H, Singer RH. 2021. Cellular variability of nonsense-mediated mRNA decay. Nature communications 12: 7203.

Sheth U, Parker R. 2006. Targeting of aberrant mRNAs to cytoplasmic processing bodies. Cell 125: 1095–1109.

Singh AK, Choudhury SR, De S, Zhang J, Kissane S, Dwivedi V, Ramanathan P, Petric M, Orsini L, Hebenstreit D et al. 2019. The RNA helicase UPF1 associates with mRNAs co-transcriptionally and is required for the release of mRNAs from gene loci. eLife 8.

Stalder L, Muhlemann O. 2009. Processing bodies are not required for mammalian nonsense-mediated mRNA decay. RNA 15: 1265–1273.

Sun X, Perlick HA, Dietz HC, Maquat LE. 1998. A mutated human homologue to yeast Upf1 protein has a dominant-negative effect on the decay of nonsense-containing mRNAs in mammalian cells. Proc Natl Acad Sci U S A 95: 10009–10014.

Szczesny RJ, Kowalska K, Klosowska-Kosicka K, Chlebowski A, Owczarek EP, Warkocki Z, Kulinski TM, Adamska D, Affek K, Jedroszkowiak A et al. 2018. Versatile approach for functional analysis of human proteins and efficient stable cell line generation using FLP-mediated recombination system. PLoS One 13: e0194887.

Weng Y, Czaplinski K, Peltz SW. 1996. Genetic and biochemical characterization of mutations in the ATPase and helicase regions of the Upf1 protein. Mol Cell Biol 16: 5477–5490.

Yoine M, Ohto MA, Onai K, Mita S, Nakamura K. 2006. The lba1 mutation of UPF1 RNA helicase involved in nonsense-mediated mRNA decay causes pleiotropic phenotypic changes and altered sugar signalling in Arabidopsis. Plant J 47: 49–62.

