## Supplementary Figures 1-4 for "UPF1 shuttles between nucleus and cytoplasm independently of its RNA binding and ATPase activities"

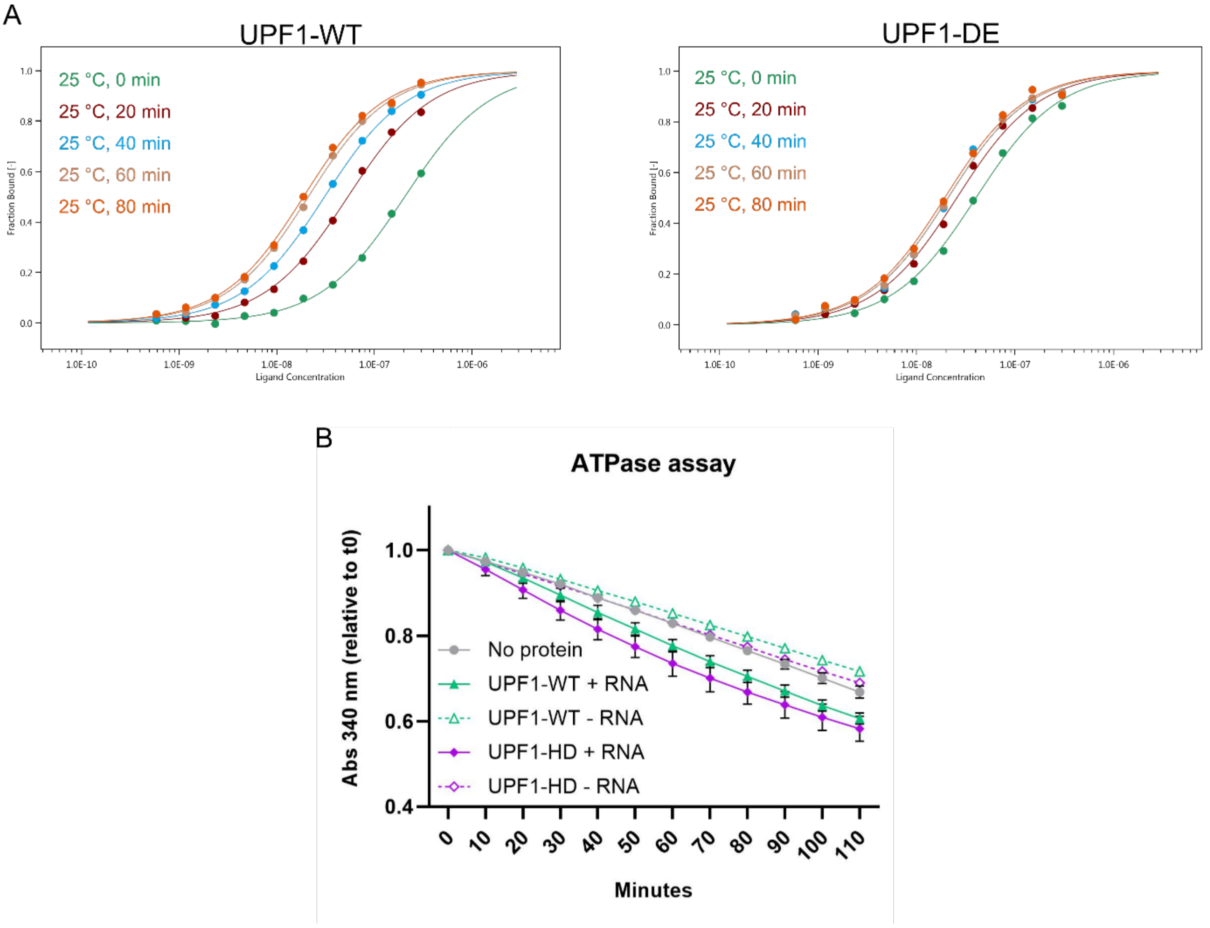


**Supplementary Figure 1**. (A) Assessment of the required time to reach the binding equilibrium for the UPF1-WT and UPF1-DE recombinant proteins. Binding reactions were incubated at 25 °C for 0, 20, 40, 60, or 80 minutes, followed by MST. A single measurement is shown. (B) NADH-coupled ATPase assay was performed to compare the ATPase activities of the full-length UPF1-WT and the ATPase/helicase domain of UPF1 (UPF1-HD) in the presence or absence of poly(U) RNA. Shown are the averages and standard deviations of 2 independent measurements.

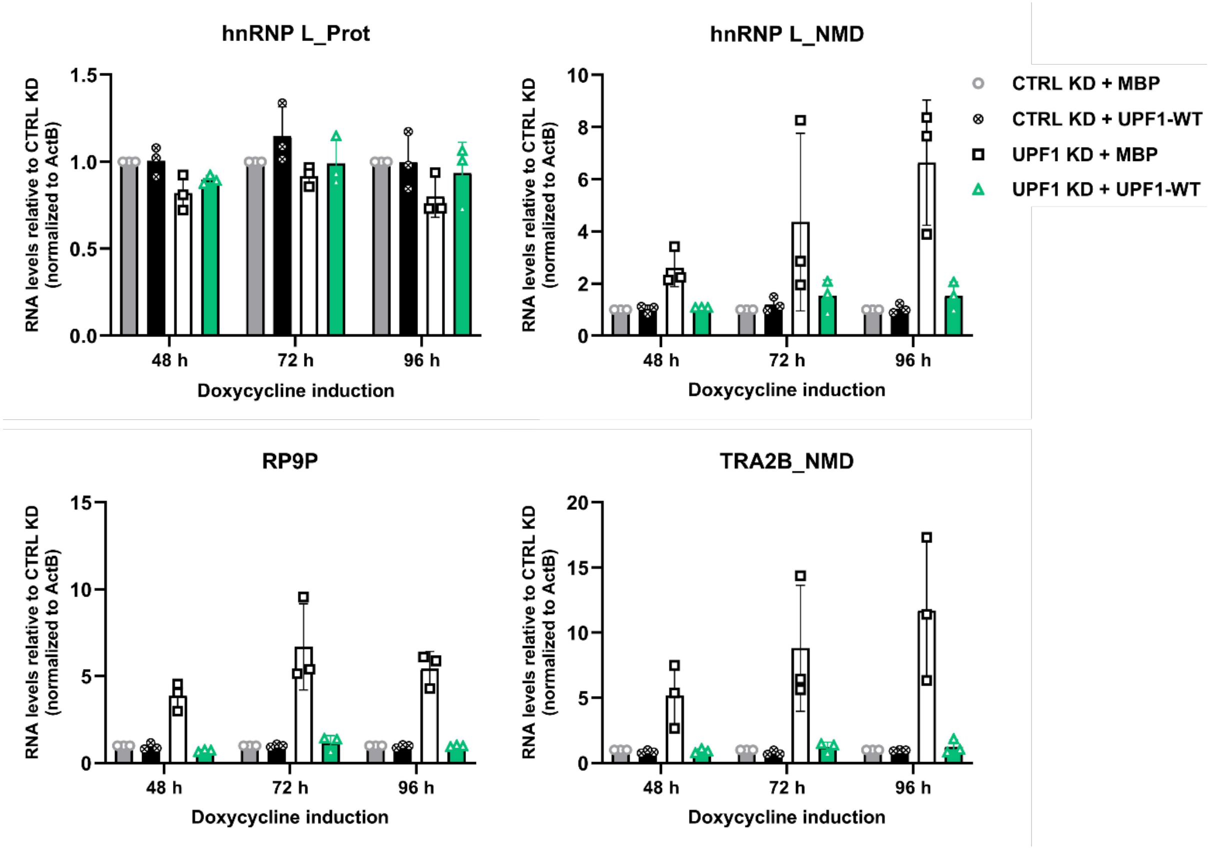


**Supplementary Figure 2**. Time-course evaluation of the NMD inhibition achieved by Doxycycline induction of HEK cells stably transfected with different pKK plasmids. Relative mRNA levels of hnRNPL_Protein, hnRNPL_NMD, RP9P, TRA2B NMD, normalized to actin, were analyzed by RT-qPCR in HEK cells 48, 72, or 96 hours after the addition of Dox. Averages and standard deviations of three independent experiments are shown.

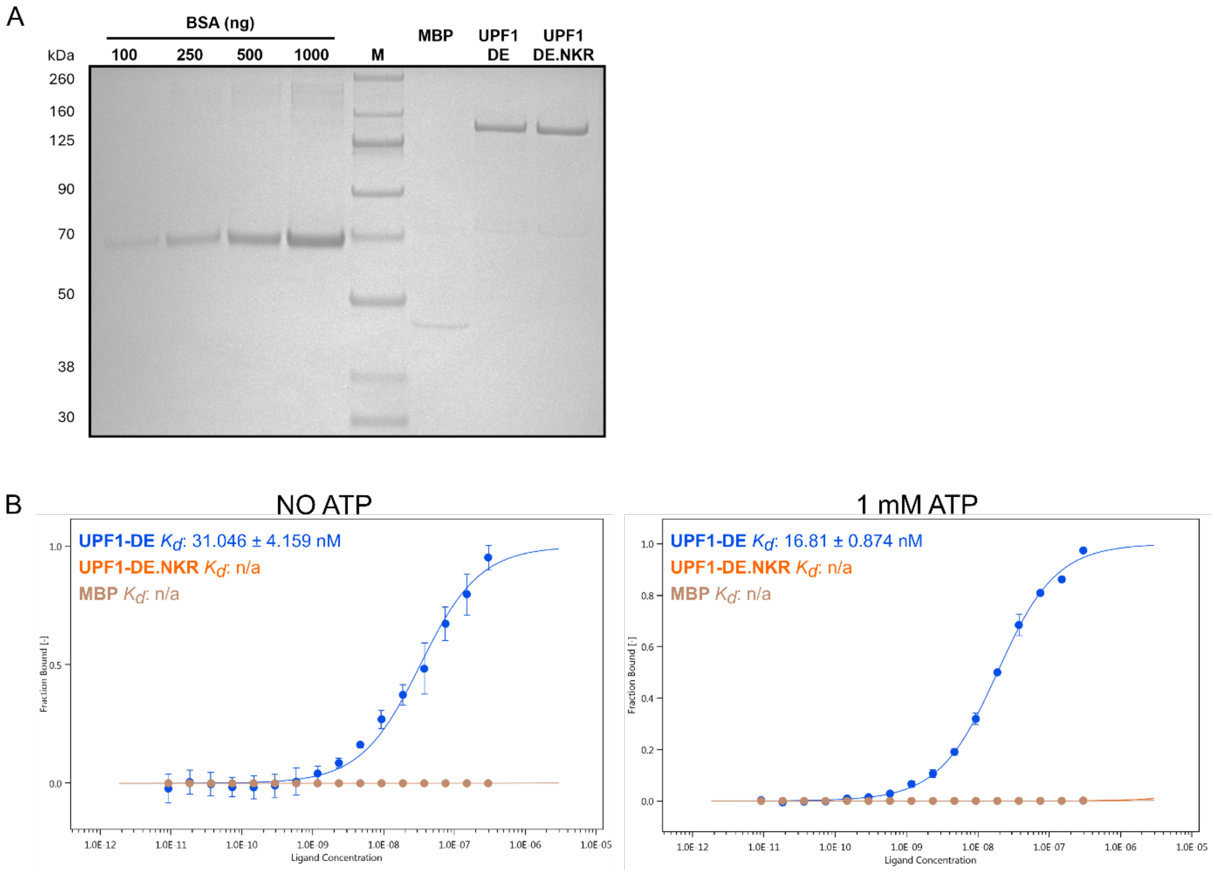


**Supplementary Figure 3**. Coomassie-stained gel showing the purified UPF1-3xFlag DE and DE.NKR mutants, and MBP-3xFlag as a control. A BSA standard curve on the same gel was used to estimate the concentration of the purified proteins. M = molecular weight marker. (B) *In vitro* RNA binding was analyzed by MicroScale Thermophoresis (MST). Binding reactions were incubated for 1 h at 25 °C in the absence or presence of ATP, followed by MST. The fraction of bound RNA is plotted against the final concentration of the unlabeled titrated proteins, and the averages and standard deviations from two independent dilution series of UPF1 are shown. Estimated dissociation constants ($K_d$) are shown for the UPF1-DE protein; n/a = not available.

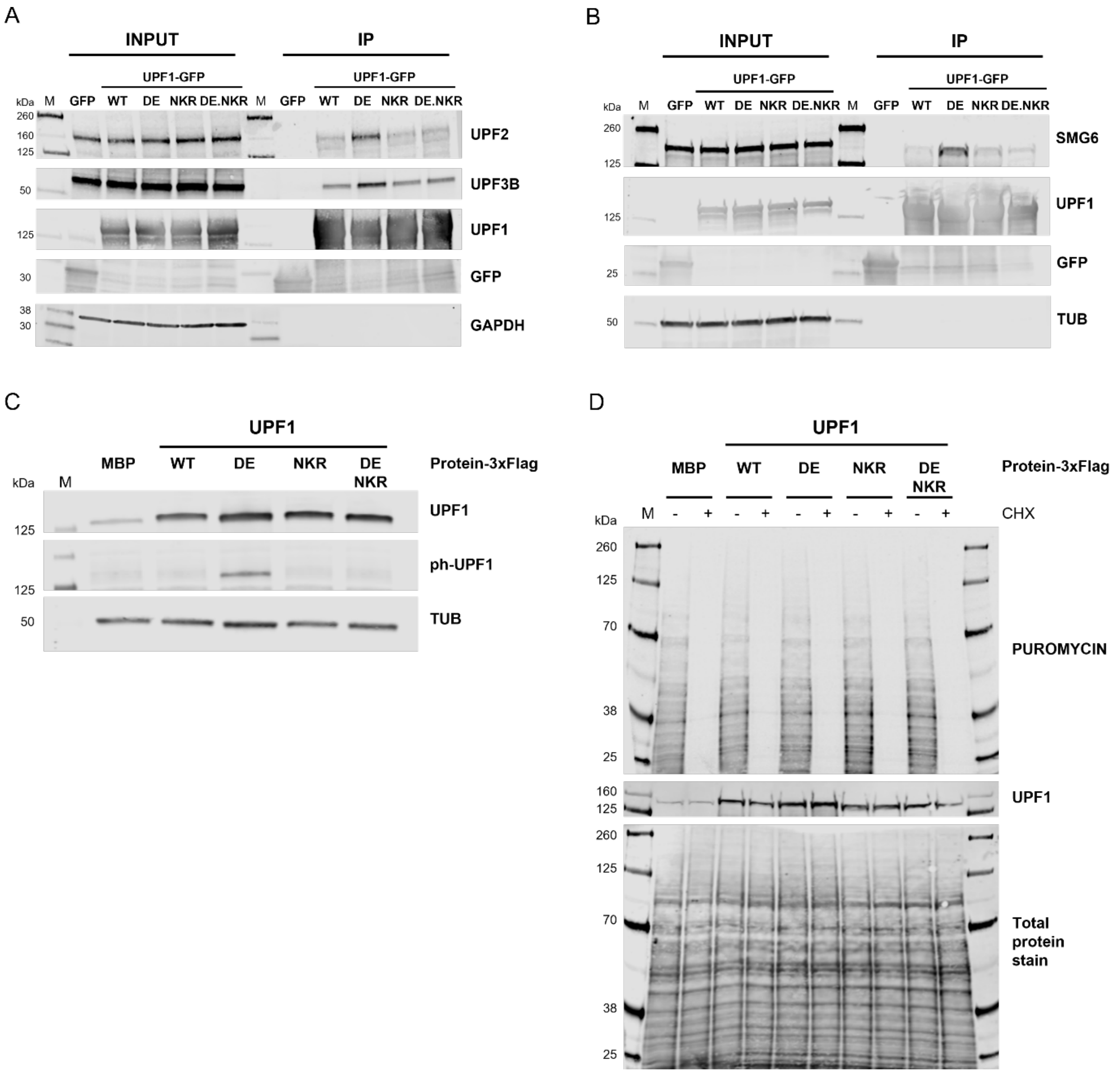


**Supplementary Figure 4**. (A-B) Western Blotting to analyze co-immunoprecipitation experiments performed with anti-GFP beads in HEK cells expressing UPF1-GFP fusions (WT, DE, NKR, or DE.NKR mutants), or GFP alone as a control. 1.5 % of the starting material (Input) and 50% (in A) or 100% (in B) of the immunoprecipitated material were loaded on an SDS-PAGE. The membranes were probed against UPF1, UPF2, UPF3B, SMG6, and GFP. GAPDH and TUB were used as controls. (C) Western blotting to assess the levels of phosphorylated UPF1 after 48 h of expression of the indicated proteins. TUB was used as a loading control. (D) Puromycin incorporation assay to assess the translation activity in HEK cells after 48 h of expression of the indicated proteins. UPF1 expression levels and puromycin incorporation in newly translated proteins were analyzed by western blotting using the indicated antibodies. Total protein stain served as loading control. Every culture was treated with cycloheximide (CHX) for 2 h before the puromycin pulse, as a control. M = molecular weight marker.
